# COGNITIVE HETEROGENEITY AND AGING: AN AGE-INDEPENDENT CILIARY GENE PROGRAM TRACKS COGNITIVE RESILIENCE IN MICE

**DOI:** 10.64898/2026.10.23.761367

**Authors:** S. Logan, A. Unnikrishnan, M.P. Baier, K. Kurup, D. Drevets, C.M. Diaz-Garcia, W. Jung, T.G. Oh, K. Ding, W.E. Sonntag

## Abstract

Cognitive decline with age is often viewed as inevitable, yet older animals show striking heterogeneity in functional trajectories, with many preserving cognitive integrity despite physiological decline. The biological basis of this resilience remains poorly understood, partly due to insensitive behavioral testing paradigms. Using an automated home-cage platform and hippocampal RNA sequencing, we integrated behavioral phenotyping with transcriptomic profiling to disentangle molecular programs linked to cognitive performance from chronological age. Behavioral data from 906 C57BL/6 mice across 129 standardized studies, trajectory-based modeling, variance-partitioned correlation, and stability-driven gene prioritization identified a transcriptional signature (271 genes) linked to cognitive state independent of age. Preserved cognition was associated with higher expression of genes governing ciliary structure and motility, neurite organization, and synaptic architecture, while impairment was associated with neuroimmune and metabolic-stress programs. A 38-gene cilia structural module score remained associated with resilience after controlling for age (partial rho = −0.81), with consistent effects across leave-one-out resampling. Because the constituent genes were selected on this same outcome, the module-level magnitude is not an unbiased effect-size estimate; recalibration against a null that reapplies the full selection procedure gave an empirical p of 0.026. This divergence emerged among genetically identical mice, indicating non-genetic, discrete hippocampal molecular signatures underlying individual differences. Cell-type attribution against the Allen Brain Cell Atlas localized the module predominantly to ependymal cells, with 27 of 38 genes ependymal-restricted (median detection 90.2% in ependymal cells versus 1.1% in CA1 glutamatergic neurons) and three genes (Dnah9, Dnai4, Kif9) also detected in CA1. The association was not attributable to ependymal tissue content: the module score was unrelated to Foxj1 expression (rho = −0.016) and the partial correlation was unchanged after adjustment for ependymal and choroid plexus marker panels, indicating that the module reflects how strongly the ciliary program is expressed rather than how many ependymal cells each sample contained. These findings support a conceptual shift: cognitive trajectories appear shaped by intrinsic molecular set points rather than by chronological age, with expression of a motile ciliary program at the cerebrospinal fluid interface, rather than the abundance of the cells carrying it, emerging as a candidate correlate of resilience. This approach offers a generalizable strategy for dissecting functional heterogeneity from aging and identifying pathways for hypothesis-driven tests of resilience across the lifespan.

## INTRODUCTION

Age-related cognitive impairment represents a major scientific and public health challenge, yet the underlying cellular and molecular mechanisms remain poorly understood^1^. A key concept emerging across species is that cognitive aging is not uniform. Substantial inter-individual variability is observed in humans^2^, non-human primates^3^, and rodents^4^, with some individuals maintaining performance comparable to young subjects while others exhibit marked decline. These observations have led to the concepts of cognitive reserve and resilience, which describe mechanisms that preserve function despite aging or pathology^5–7^. Notably, such heterogeneity extends beyond normative aging to neurodegenerative disorders, including Alzheimer’s and Parkinson’s disease, where cognitive outcomes often dissociate from pathological burden^7–11^. Defining the biological mechanisms that underlie this variability is therefore essential for understanding both normal brain aging and cognitive vulnerability across disease states^12,13^. Critically, the determinants of these divergent trajectories need not arise during aging itself^14–20^: if cognitive resilience and vulnerability are established earlier in life, studying animals only at advanced ages may capture the downstream consequences of trajectories that were already established, rather than their origin.

Despite this need, current experimental approaches in rodent models, the primary system for mechanistic studies, have been limited in their ability to resolve cognitive heterogeneity. Most studies rely on cross-sectional comparisons using small cohorts and behavioral assays that are poorly suited to detect inter-individual variability within age groups. These limitations obscure biologically meaningful variation and restrict identification of mechanisms underlying resilience^21^. In addition, inconsistencies in testing conditions, strain differences, and experimenter-dependent variability further reduce reproducibility^22,23^. Together, these factors have contributed to a broader lack of confidence in preclinical behavioral measures of aging^24^ (**Supplemental Tables 1 and 2**), reinforcing the assumption that cognitive decline is an inevitable consequence of aging rather than a heterogeneous and potentially modifiable process.

Automated behavioral platforms provide a strategy to overcome these limitations by enabling standardized, high-throughput assessment of cognitive function with minimal experimenter interference^25–34^. However, their application to aging remains limited, and their capacity to resolve individual cognitive trajectories across the lifespan has not been systematically evaluated. Moving beyond categorical comparisons toward continuous measures of cognitive performance may substantially improve the resolution of experimental studies, particularly in heterogeneous conditions such as aging and neurodegeneration. Integrating high-dimensional behavioral phenotyping with trajectory-based modeling and variance-partitioned molecular analyses has the potential to provide a framework for identifying mechanistically relevant pathways linked to cognitive outcomes.

Here, we performed a comprehensive analysis of cognitive function using 129 mouse aging studies comprising 906 animals of both sexes, tested under standardized automated conditions over a five-year period. We sought to determine whether reproducible behavioral measures can resolve cognitive heterogeneity and define individual trajectories independent of chronological age. We show that a subset of behavioral metrics provides stable and biologically sensitive measures of cognition, reveals early-emerging heterogeneity, and supports trajectory-based classification of cognitive states. Integration with hippocampal gene expression identifies transcriptional programs associated with cognitive performance that are largely separable from age-dependent effects. These programs are enriched for pathways related to ciliary structure and motility, neurite organization, and synaptic structure, implicating cilium-associated processes as a key biological axis underlying cognitive heterogeneity. Although not focused on specific disease models, this framework provides a generalizable approach for separating cognitive outcomes from aging or disease processes and for identifying mechanisms of vulnerability and resilience across heterogeneous neurological conditions.

## RESULTS

### High-dimensional cognitive phenotyping identifies stable behavioral metrics across years

A major barrier to standardizing measures of cognitive aging is limited reproducibility across time, cohorts, and experimental conditions. To identify behavioral metrics that are both stable and biologically informative, we analyzed a large multi-year dataset of 906 C57BL/6 mice drawn from 129 studies conducted between 2020 and 2024 under standardized automated conditions (**Figure 1a–d**). Thirteen cognitive variables (**Supplemental Table 3a**) and five movement parameters (**Supplemental Tables 4a and 4b**) were initially evaluated.

**Figure 1:**
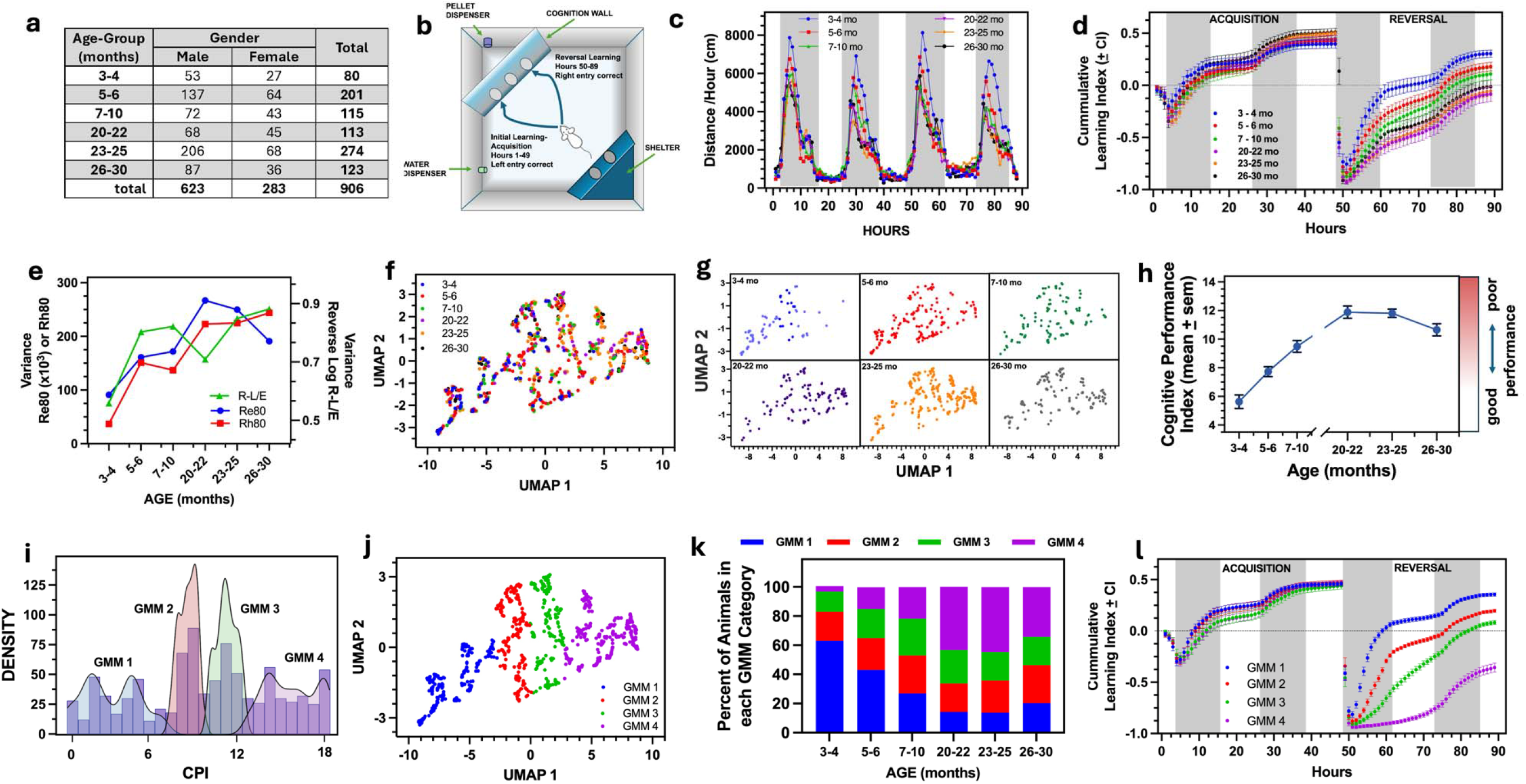
Behavioral characterization and identification of cognitive heterogeneity across age. (A) Cohort selection. A total of 906 C57BL/6 mice were selected from a database of >3,400 animals (129 studies) tested between 2020 and 2024 under identical behavioral conditions. Only non-manipulated control animals were included. Animals failing to reach ≥70% success during initial training were excluded. **(B)** Schematic of the automated behavioral phenotyper. During the acquisition phase (0–49 h), animals learned to enter the left side for a pellet reward (FR5 schedule). During reversal (50–89 h), the correct side was switched to the right. **(C)** Distance traveled (cm/h) across the 89-h testing period by age group. Shaded regions indicate the dark phase of the L:D cycle. **(D)** Cumulative learning index across age groups (mean ± 95% CI). Acquisition learning was preserved across ages, whereas reversal learning declined progressively with age. **(E)** Variance of three key behavioral metrics (R-L/E, Re80, Rh80) increased with age, indicating greater heterogeneity among older animals. **(F)** UMAP × UMAP embedding of behavioral profiles across the lifespan. Each point represents an individual mouse colored by age group. The landscape reveals a continuous trajectory with increasing divergence in cognitive performance with age. **(G)** UMAP embeddings (from (F)) shown separately for each age group. **(H)** Mean CPI values across age groups derived from principal-curve analysis of the behavioral UMAP embedding (mean ± s.e.m.). Cognitive performance Index (CPI) increased progressively with age, indicating a shift along the behavioral trajectory toward impaired performance. **(I)** Distribution of CPI values with Gaussian mixture model (GMM) fit, revealing multiple behavioral subpopulations. Model selection supported a four-component solution. **(J)** Spatial distribution of GMM clusters within the behavioral embedding, demonstrating coherent separation of behavioral states. **(K)** Age-dependent changes in GMM composition. The prevalence of early-life GMM clusters (GMM1) decreased with age, whereas late-life clusters (GMM4) increased, indicating a progressive shift in behavioral state distribution. **(L)** Cumulative learning index stratified by GMM cluster membership (mean ± 95% CI). Stratification by behavioral state produced stronger separation of performance than chronological age alone, indicating that cognitive state is a more robust determinant of learning ability than age. Together, these analyses demonstrate that aging is associated with increasing behavioral heterogeneity and the emergence of distinct cognitive states that are independent of chronological age.

Reversal-learning measures exhibited substantially lower between-year variance than acquisition metrics, indicating greater longitudinal stability. Linear mixed-effects modeling^35–37^ showed that variance attributable to ‘Year’ accounted for 4.6–8.8% of total variance during Acquisition but only 0–4.4% during Reversal Learning (**Supplementary Table 3b**). Variance associated with combined Year + Experiment was consistently lower for reversal variables, demonstrating resistance to procedural variability. Robust main effects of Age were observed across all reversal metrics (p<0.001), whereas acquisition measures showed more variable age sensitivity. Significant Sex effects were detected for most reversal measures, with females reaching reversal criteria faster than males, although no Age × Sex interactions were observed.

Among these metrics, R-L/E (capturing adaptive behavioral flexibility during early reversal learning) combined low procedural variance (∼2.5%) with strong Age and Sex effects (p<0.01), highlighting its interpretive value. Collectively, these findings identify reversal learning as a stable and biologically sensitive behavioral domain for studying cognitive aging.

### Cognitive aging is characterized by increasing behavioral heterogeneity organized along a continuous performance axis

Having established that the behavioral measures are reproducible over time, we next examined whether these metrics could resolve inter-individual variability in cognitive aging and define the structure of behavioral heterogeneity across the lifespan. Analyses were restricted to inbred animals to minimize genetic variability, allowing observed differences to primarily reflect endogenous aging processes.

Behavioral variance increased progressively with age (**Figure 1e**), emerging as early as 5–6 months and reaching maximal levels at 20–22 months (p < 0.0001). Reversal-learning measures provided the most sensitive readout of this heterogeneity. Notably, substantial variability was observed within age groups, indicating that chronological age alone does not fully determine cognitive ability.

Principal component analysis of three key reversal-learning metrics (Re80, Rh80, and R-L/E) revealed a dominant shared dimension of performance. The first principal component accounted for 84.7% of total variance and exhibited concordant loadings across measures, indicating a common behavioral axis. Age groups occupied overlapping regions of this space, further supporting the presence of within-age heterogeneity rather than discrete age-defined states.

To visualize this structure, nonlinear dimensionality reduction using UMAP was applied to the multivariate behavioral data (**Figure 1f, g**). A principal curve fitted to the UMAP embedding generated a continuous measure of cognitive performance across individuals. Each animal’s position along this curve was defined as the Cognitive Performance Index (CPI), representing an ordered, continuous measure of cognitive state derived from behavioral performance rather than chronological age. CPI values (**Figure 1h**) were strongly correlated with the underlying behavioral metrics (r=0.69–0.88), supporting its validity as an integrative measure of cognitive performance.

Gaussian mixture modeling (GMM) of the CPI distribution identified discrete behavioral subgroups embedded within this continuous axis (**Figure 1i, j; Supplemental Figure 1,2**). These subpopulations represent progressively impaired performance states and extended beyond traditional binary classifications. The relative abundance of these states shifted systematically with age, with early-life states decreasing and late-life states increasing in prevalence, indicating redistribution of animals along the cognitive performance axis rather than uniform decline.

Independent validation using GMM values confirmed progressive reductions in performance across CPI-defined subgroups during reversal learning (**Figure 1k**). Importantly, unsupervised stratification by CPI-defined state produced markedly greater separation in performance than chronological age alone, demonstrating that cognitive state is a more precise determinant of behavioral outcome (**Figure 1l**).

Together, these analyses demonstrate that cognitive aging is characterized by increasing behavioral heterogeneity organized along a continuous performance axis with embedded discrete states, and that cognitive performance is only partially explained by chronological age.

### Disentangling molecular mechanisms of aging from cognitive performance reveals structured programs

The behavioral framework described above enables direct separation of molecular processes associated with chronological aging from those linked to cognitive performance. To define how behavioral variation maps onto the underlying biology, hippocampal RNA-seq data were integrated with CPI and age for each animal (**Figure 2a–d**).

**Figure 2.**
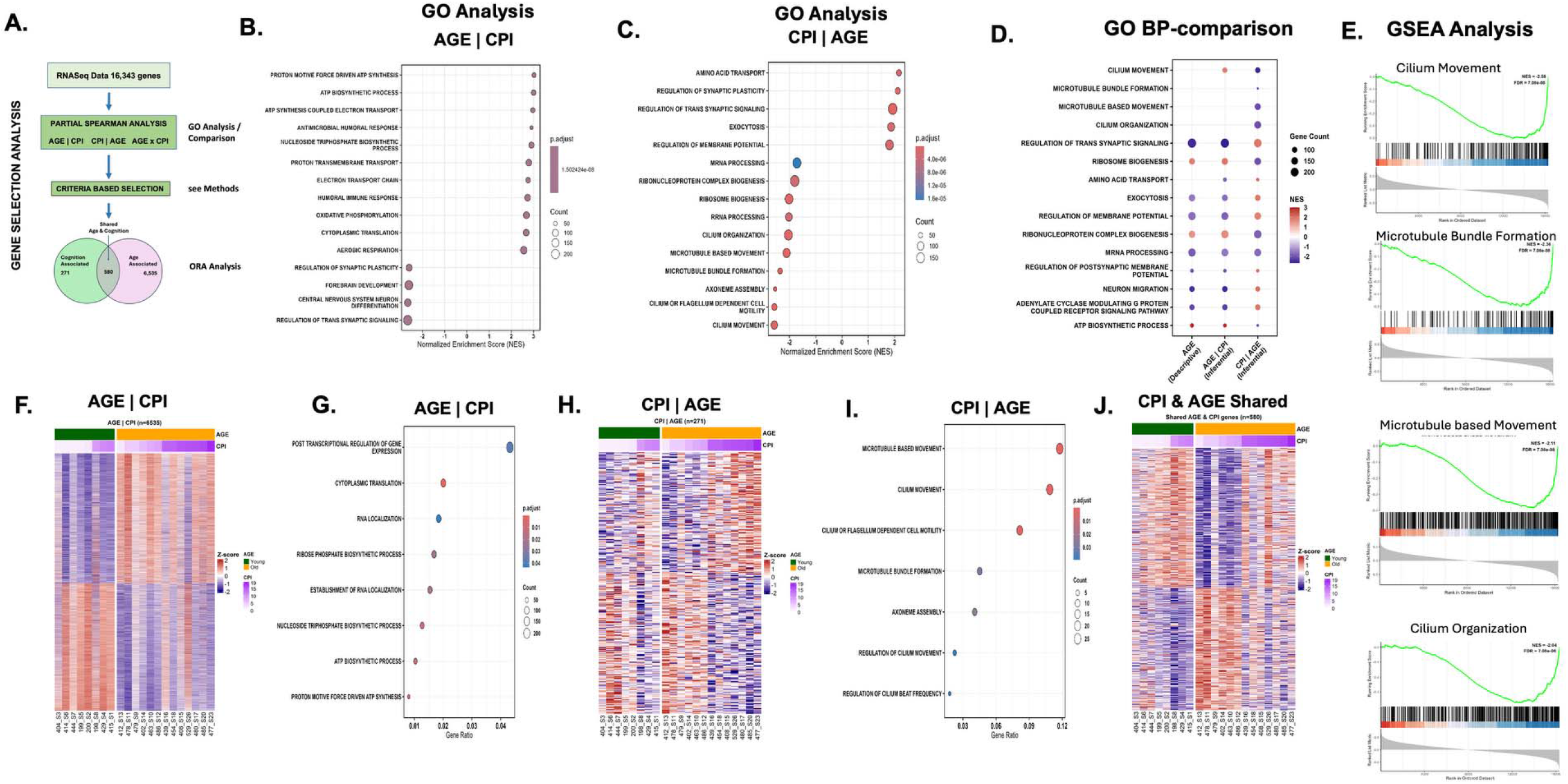
Separation of age- and cognition-associated transcriptional programs reveals distinct molecular pathways linked to cognitive performance. **(A)** Analytical framework for partitioning gene expression into components associated with chronological age and cognitive performance index (CPI). Gene–trait relationships were evaluated using partial Spearman correlations to define age-associated (Age|CPI), cognition-associated (CPI|Age), and shared (Age×CPI) transcriptional programs. **(B,C)** Gene ontology (GO Biological Process) enrichment analysis of all detected genes (n = 16,343) identified distinct functional profiles. Age-associated genes (Age|CPI) were enriched for metabolic, cellular, and immune-related processes, whereas CPI-associated genes (CPI|Age) were enriched for neuronal signaling, synaptic plasticity, and cytoskeletal pathways, including ciliary and microtubule-related processes. **(D)** Comparative GO analysis using descriptive-analysis based gene assignments (Age), and inferential analysis (Age|CPI, and CPI|Age) gene sets highlight the emergence of cilia- and microtubule-associated pathways specifically within cognition-associated programs after accounting for chronological age. **(E)** Gene set enrichment analysis (GSEA) demonstrates that pathways related to ciliary movement, microtubule bundle formation, microtubule-based movement, and cilium organization are significantly enriched along the ranked gene list, indicating a non-random distribution of these pathways in relation to cognitive performance. **(F–J)** Heatmap visualization and over-representation analysis (ORA) of gene sets defined by partial Spearman classification further resolve functional distinctions between Age|CPI (F,G), CPI|Age (H,I), and shared (J) transcriptional programs. CPI-associated genes (n = 271) show strong enrichment for cilia- and microtubule-based processes, whereas age-associated genes (n = 6,535) are dominated by metabolic and systemic pathways. The AgexCPI gene set (580 genes) did not converge on a specific biological process that reached significance and is not shown, Notably, although the majority of transcriptional variation was associated with chronological age (n = 6,535 genes), a much smaller CPI-associated gene set (n = 271 genes) exhibited strong and specific enrichment for ciliary and microtubule pathways, indicating that cognitive performance is linked to a focused and functionally coherent molecular program not explained by age alone. These cognition-associated programs support the existence of trajectory-specific molecular mechanisms underlying cognitive heterogeneity. Notably, these cognition-associated pathways are consistent with early-emerging, trajectory-defining biological processes that are not fully captured by chronological age.

As an initial step, conventional bivariate (descriptive) Spearman correlation analyses were used to classify genes associated with age. This approach identified transcripts correlated with cognitive performance and age (**Supplemental Figure 3** and data not shown). However, because performance and age are correlated, descriptive analyses cannot distinguish whether the gene expression changes reflect chronological aging, cognitive state, or their covariance.

To resolve this limitation, we applied a partial Spearman analysis that partitions shared variance between CPI and Age based on several criteria (**Supplemental Table 5**). This approach enabled classification of genes associated with cognitive performance independent of age (CPI | Age), genes associated with chronological aging independent of cognitive performance (Age | CPI), and genes reflecting shared variance (CPI × Age) (**Figure 2a–c**).

Comparison of descriptive and inferential analyses revealed a marked asymmetry in gene associations (**Figure 2d**). While the majority of genes were associated with chronological age (n = 6,535), a substantially smaller subset was associated with cognitive performance independent of age (n = 271), with an additional intermediate set reflecting shared variance (n = 580).

To assess whether the number of genes identified by partial Spearman correlation exceeded chance expectation, we performed permutation-based empirical false discovery rate (FDR) estimation. Cognitive Performance Index (CPI) values were randomly shuffled across animals while gene expression data were held fixed, and the number of genes meeting the |r| ≥ 0.44 threshold was recorded across 1,000 permutations. The observed gene count (n = 271) was approximately 18.6-fold greater than the permutation mean (14.5 genes; range 0–94), placing the observed result entirely outside the permutation null distribution and yielding an empirical FDR of 5.4% (**Supplemental Figure 4**). This result confirms that the identified cognition-associated gene set reflects a structured biological signal rather than a consequence of multiple comparisons at this sample size.

The empirical FDR of 5.4% marginally exceeds the conventional 5% threshold but represents a conservative upper bound on false discovery for the full candidate set. To further prioritize genes with stable, reproducible associations, we applied a Cognitive Stability Index (see Methods) as a continuous ranking metric rather than a binary filter. The CSI integrates effect size with leave-one-out (LOO) resampling stability, specifically sign consistency and variance of LOO estimates across individuals, such that spurious correlations arising by chance are systematically depressed in the ranking because random associations do not survive resampling based on consistent direction and low variance. This two-stage framework, in which a permutation-validated effect-size threshold (|r| ≥ 0.44) defines the candidate set and CSI-based stability ranking deprioritizes residual false positives, provides more stringent false discovery control than either criterion applied in isolation, without imposing an arbitrary secondary cutoff. Together, these results indicate that cognitive performance is not driven by global transcriptional changes but instead reflects a selective and structured molecular program.

### Functional enrichment analyses reveal distinct molecular programs associated with cognitive performance and aging

Functional enrichment analyses were performed to define the biological pathways associated with each transcriptional category (**Figure 2b–j**). Age-associated genes (Age | CPI) were enriched for metabolic, mitochondrial, and immune-related pathways, consistent with established hallmarks of biological aging^38,39^ (**Supplemental Figure 3**). These pathways were broadly distributed and reflected systemic physiological processes, including energy metabolism, oxidative stress responses, and inflammatory signaling.

In contrast, genes associated with cognitive performance independent of age (CPI | Age) (**Figure 2c**) were strongly enriched for pathways related to cilium organization, ciliary movement, microtubule-based processes, intracellular transport, and neuronal signaling. These enrichments were highly specific and not prominent among age-associated genes, indicating that transcriptional variation linked to cognitive performance reflects a focused and functionally coherent molecular program rather than generalized aging physiology. Gene set enrichment analysis (**Figure 2e**) confirmed that these pathways were non-randomly distributed along the ranked gene list, with ciliary and microtubule-related processes consistently enriched in association with cognitive performance.

Redundancy analysis demonstrated that enrichment of cilia-related pathways was not driven by repeated annotation of a limited number of genes but instead reflected a distributed transcriptional module spanning multiple pathways (**Supplemental Figures 5a-d and 6a,b**). This supports the interpretation that cilia- and microtubule-associated processes represent coordinated cellular systems linked to cognitive state.

Genes associated with both age and cognitive performance (CPI × Age) were enriched for immune signaling, metabolic and mitochondrial processes, and stress-response pathways, representing an interface through which systemic processes such as aging and infections influence cognitive function. The presence of cilia-related genes within both cognition-specific and shared categories suggests that ciliary pathways are operative at multiple levels within the age–cognition axis.

Collectively, these findings reveal a fundamental asymmetry in molecular organization: While chronological aging is associated with broad and diffuse transcriptional changes, cognitive performance is linked to a smaller, highly structured, and biologically coherent gene program dominated by ciliary and microtubule-related processes that emerge early during the lifespan.

### Stability-based classification reveals opposing molecular programs of cognitive resilience and impairment

To define molecular programs associated with preserved versus impaired cognitive performance, genes were classified based on both the direction and stability of their association with CPI. Gene–cognition relationships were first defined using partial Spearman correlation (CPI | Age), and a Cognitive Stability Index (CSI) was applied to prioritize genes exhibiting consistent associations across individuals rather than outlier-driven effects (**Figure 3a**).

**Figure 3.**
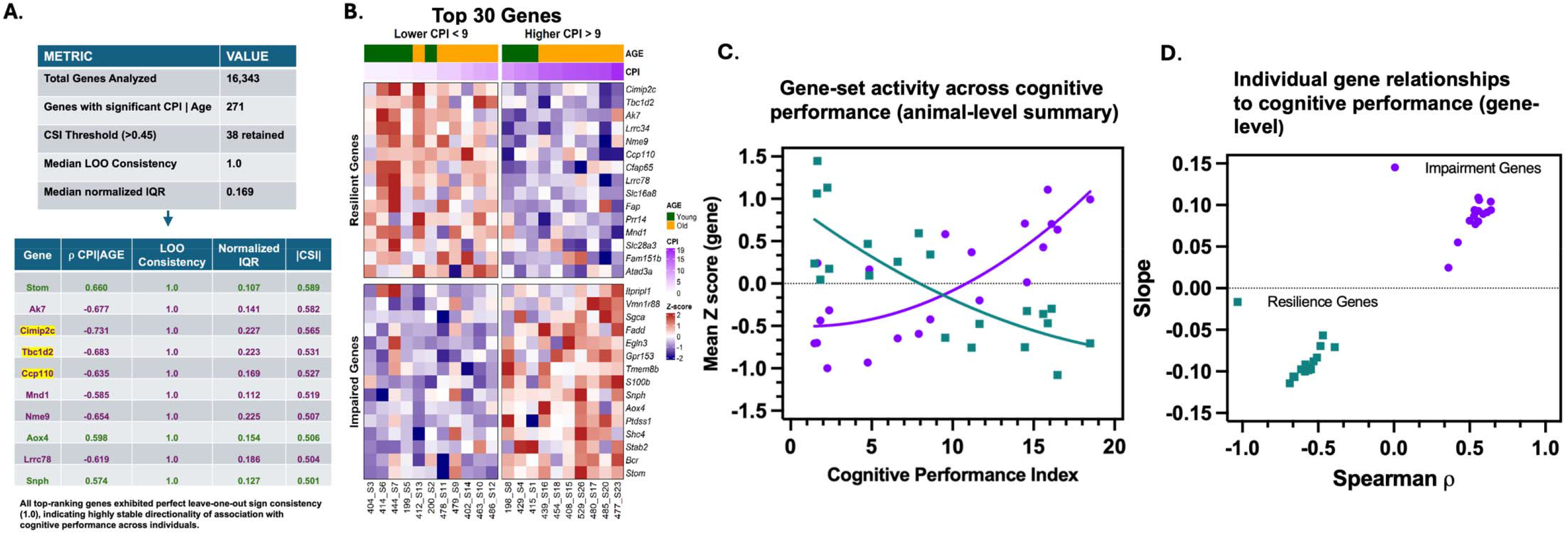
Stability-based identification of cognition-associated gene programs reveals opposing molecular signatures of resilience and impairment. (A) Gene–cognition relationships were evaluated using partial Spearman correlation analysis (CPI | Age) to isolate associations with cognitive performance independent of chronological age. To prioritize robust and reproducible signals, a Cognitive Stability Index (CSI) was calculated for each gene by integrating the magnitude of the partial Spearman correlation, leave-one-out (LOO) sign consistency, and variability of LOO estimates (normalized IQR). This framework identifies genes whose associations with cognitive performance are both strong and stable across individuals. The CSI was applied as a continuous ranking metric rather than as a binary filter, so that genes with strong but unstable associations are deprioritized without imposing a secondary cutoff; candidate genes were defined by the permutation-validated effect-size threshold described in the Results. (B) High-confidence cognition-associated genes were partitioned based on the direction of their association with cognitive performance. Genes with positive partial correlations (CPI | Age) were classified as resilience-associated, whereas genes with negative correlations were classified as impairment-associated. A CPI threshold (CPI > 9 vs. CPI < 9) is shown for visualization of high- and low-performing animals and does not influence gene selection. This classification defines two opposing molecular programs linked to cognitive state. (C) Gene-set activity across cognitive performance at the animal level. Each point represents an individual animal plotted by cognitive performance index (CPI) and the mean standardized expression (z-score) of genes within each gene set. Gene expression values were z-scored across animals and averaged within each animal for resilience- or impairment-associated gene sets. Locally weighted regression (loess) curves illustrate trends across the cognitive spectrum, with resilience-associated genes (green) increasing and impairment-associated genes (purple) decreasing with improved cognitive performance. (D) Gene-level relationships with cognitive performance. Each point represents an individual gene plotted by its Spearman correlation (ρ) with CPI (x-axis) and the slope of the linear relationship with CPI (y-axis), reflecting both direction and magnitude of association. Gene expression values were standardized across animals. Resilience-associated genes (green) exhibit positive associations with cognitive performance, whereas impairment-associated genes (purple) exhibit negative associations, demonstrating a bidirectional molecular organization aligned with cognitive state. Collectively, these analyses demonstrate that cognition-associated molecular programs are defined by stable and reproducible relationships with cognitive performance across individuals, rather than by large but inconsistent expression differences, enabling the identification of biologically coherent gene sets underlying cognitive resilience and impairment.

High-confidence cognition-associated genes were partitioned based on the direction of their association with CPI (**Figure 3b**). Genes positively associated with cognitive performance were classified as resilience-associated, whereas genes negatively associated with cognitive performance were classified as impairment-associated, defining two opposing molecular programs aligned with cognitive state (**Figure 3b–d**).

Resilience-associated genes were strongly enriched for ciliary and centrosomal components, including regulators of ciliogenesis, axonemal assembly, and intracellular transport, implicating cilia-dependent signaling and structural organization as key determinants of preserved cognitive function. In contrast, impairment-associated genes were enriched for immune and neuromodulatory pathways, consistent with neuroimmune activation and altered signaling contributing to functional decline. Genes shared between aging and cognitive performance were enriched for mitochondrial and metabolic pathways, suggesting convergence between systemic aging processes and cognitive vulnerability.

Together, these findings demonstrate that cognitive decline reflects divergence along distinct molecular trajectories defined by stable and reproducible gene expression programs, rather than uniform age-dependent deterioration. The consistent enrichment of ciliary and microtubule-related processes within cognition-associated and resilience-linked gene sets identifies these pathways as candidate mechanisms underlying cognitive heterogeneity during aging.

### Statistical Sensitivity of Gene-Behavior Correlations

A sensitivity analysis was conducted to characterize the statistical properties of the gene-CPI correlation approach. For n = 21 with a two-tailed alpha of 0.05, the minimum detectable effect size at 80% power is approximately r ≈ 0.55–0.58. The applied threshold of |r| ≥ 0.44 in this study, therefore, captures an exploratory range that includes correlations below the conventionally well-powered threshold. Correlations exceeding |r| ≥ 0.55 within this dataset are reliably powered, while those in the range 0.44–0.55 should be considered preliminary and prioritized for follow-up validation. Additionally, the all-male composition of the RNA-seq cohort precludes sex as a confound in gene-behavior correlations within this dataset. Although sex effects on reversal learning performance are reported in the broader behavioral analyses, these effects cannot confound transcriptomic-behavioral associations in a single-sex sample. The relevance of sex as a biological variable in the transcriptomic findings would need to be addressed in future studies incorporating female animals.

### Cilia structural gene expression is reduced in cognitively impaired animals

The CPI|Age gene set is functionally heterogeneous, comprising cilia structural and motility genes as well as immune, metabolic, and other transcripts. To specifically test the relationship between cilia structural gene expression and cognitive performance, we computed a module score representing the mean VST-normalized expression of the 38 bona fide cilia structural genes within the 271 gene CPI|Age set across 21 animals. The cilia module score ranged from 7.27 to 7.998 (mean 7.58, SD 0.20) across animals.

Cilia module scores were significantly higher in cognitively intact animals (mean 7.72) compared with cognitively impaired animals (mean 7.43; Welch t-test: t = 5.22, df = 17, p = 0.0001), defined by a median CPI split (n = 11 intact, n = 10 impaired). Age alone did not explain this difference: young (5-6 month) and old (21-25 month) animals had nearly identical cilia module scores (7.62 vs. 7.56; t = 0.62, df = 10.1, p = 0.55), indicating that higher cilia structural gene expression was associated with preserved cognition rather than chronological age.

Partial Spearman correlation analysis, controlling for age, confirmed a strong negative association between the cilia module score and CPI (partial rho = −0.81, p < 0.001), indicating that animals with higher cilia structural gene expression performed markedly better on cognitive testing. Unlike the heterogeneous full gene set, the unadjusted correlation was also strong and highly significant (rho = −0.72, p = 0.0002), and controlling for age further strengthened the association, consistent with age acting as a partial suppressor of the underlying cognitive relationship. Restricting the module score to bona fide cilia structural genes thus produced a substantially stronger and more robust association with cognition than the full 271-gene set, identifying cilia structural gene expression as a focused correlate of cognitive resilience.

### Cilia structural genes are uniformly and stably associated with cognitive resilience

We next examined whether the resilience association of cilia structural genes was uniform across the gene set and robust to individual animals. For each of the 38 cilia structural genes, we computed the partial Spearman correlation with CPI (controlling for age) and a Cognitive Stability Index (CSI) derived from leave-one-out resampling, which integrates effect size, sign consistency, and dispersion across resampling iterations (**Figure 3**).

Across all 38 cilia structural genes, the direction of the CPI association was remarkably consistent: 36 of 38 genes were resilience-associated (higher expression with lower CPI), with only two genes (Lrrc39, Ccdc177) showing the opposite direction. Leave-one-out resampling demonstrated perfect sign consistency (1.0) for all 38 genes, indicating that the directional association of every gene was preserved regardless of which animal was excluded. Sixteen of the 38 genes (42%) reached both FDR significance and moderate-to-high stability (CSI >= 0.3), and five genes (Daw1, Ccp110, Ak7, Dnai4, and Cimip2c) met the high-stability threshold (CSI >= 0.5), with Daw1 showing the strongest and most stable association (partial rho = −0.79, CSI = 0.79). These genes represent the highest-confidence cilia structural correlates of cognitive resilience in this dataset.

### Cell-type attribution localizes the cilia structural module to ependymal cells

To determine which hippocampal cell types express the genes of the CPI|Age set, we first queried the Allen Brain Cell Atlas cluster marker gene database. Of the 271 genes, 13 were returned as cluster-defining markers for hippocampal populations. Twelve of the 13 marked neuronal populations, and one (Ccdc146) marked an ependymal population (Ependymal NN, Astro-Epen class). None of the 38 cilia structural genes was among these 13, so this query does not address the cellular origin of the cilia module, and we therefore examined the module genes directly.

Among the hippocampus-specific neuronal markers, four genes (Stom, Apela, Lefty2, Serpina3g) marked CA1 glutamatergic neurons (CA1-ProS Glut, IT-ET Glut class). Serpina3g additionally marked CA3 glutamatergic neurons, and Ntf3 marked both CA2 (CA2-FC-IG Glut) and dentate gyrus neurons (DG-PIR Ex IMN). The remaining 258 DEGs did not appear as cluster-defining markers in the atlas, indicating that while they are expressed across hippocampal cell types, they do not uniquely define any single population. These results are summarized in **Table 1**.

**Table 1.** Allen Brain Cell Atlas cell-type attribution of marker genes within the CPI|Age set. Of 13 unique cluster-defining marker genes, 12 identified neuronal populations; among hippocampal-specific markers, CA1 glutamatergic neurons were the most consistently represented subpopulation. These 13 genes are the subset of the 271-gene CPI|Age set returned as cluster-defining markers and do not include any of the 38 cilia structural genes; attribution of the cilia module is reported separately in the text.

| Gene | Allen Brain Atlas Cell Subclass | Allen Class | Neuronal | Hippocampal Specific<sup>1</sup> |
| --- | --- | --- | --- | --- |
| *Stom* | CA1-ProS Glut | IT-ET Glut | Yes | Yes |
| *Apela* | CA1-ProS Glut | IT-ET Glut | Yes | Yes |
| *Lefty2* | CA1-ProS Glut | IT-ET Glut | Yes | Yes |
| *Serpina3g* | CA1-ProS Glut | IT-ET Glut | Yes | Yes |
| *Ntf3* | CA2-FC-IG Glut | IT-ET Glut | Yes | Yes |
| *Ccdc146* | Ependymal NN | Astro-Epen | No | No |
| *2810459M11Rik* | OB-STR-CTX Inh IMN | OB-IMN GABA | Yes | No |
| *Ttc29* | PAG Pou4f2 Glut | MB Glut | Yes | No |
| *Lepr* | PAG-SC Neurod2 Meis2 Glut | MB Glut | Yes | No |
| *Serpinb1b* | SC-PAG Lef1 Emx2 Gaba | MB GABA | Yes | No |
| *Ebf3* | SCiw Pitx2 Glut | MB Glut | Yes | No |
| *Egln3* | Vip Gaba | CTX-CGE GABA | Yes | No |
| *Sostdc1* | Vip Gaba | CTX-CGE GABA | Yes | No |

Cell subclass and class designations follow the Allen Brain Cell Atlas whole-mouse-brain taxonomy (Yao et al., 2023). Genes marking multiple clusters are shown with their hippocampal assignment where available. Glut, glutamatergic; GABA, GABAergic; IMN, immature neuron; NN, non-neuronal; MB, midbrain; CTX, cortex; OB, olfactory bulb; PAG, periaqueductal gray; SC, superior colliculus. <sup>1</sup>Hippocampal Specific indicates whether the gene is a cluster-defining marker for a hippocampal population in the Allen reference; all 13 genes are expressed in the dissected hippocampal tissue.

**Table 2.** CIBERSORTx estimated cell-type proportions and partial Spearman correlations with the 38-gene cilia structural module score and Cognitive Performance Index (controlling for age, n = 21).

| Cell Type | Mean Absolute Score | r vs Cilia Score | p (cilia)* | r vs CPI | p (CPI)* |
| --- | --- | --- | --- | --- | --- |
| Mossy-Cyp26b1 | 1.19 | -0.31 | 0.190 | +0.45* | 0.044 |
| Granule-mature | 0.32 | -0.18 | 0.443 | -0.01 | 0.966 |
| Granule-immature | 0.29 | +0.10 | 0.665 | -0.09 | 0.721 |
| GABA-Lhx6 | 0.24 | -0.31 | 0.182 | +0.15 | 0.533 |
| Astrocytes | 0.19 | -0.24 | 0.309 | +0.27 | 0.247 |
| OL | 0.06 | -0.08 | 0.749 | -0.17 | 0.463 |
| Neuroblast 1 | 0.03 | +0.06 | 0.795 | +0.19 | 0.419 |
| Endothelial | 0.01 | -0.08 | 0.748 | +0.07 | 0.784 |
| Microglia | 0.01 | -0.08 | 0.738 | -0.02 | 0.922 |

*r*, partial Spearman correlation coefficient. \* $p < 0.05$ (uncorrected). CPI, Cognitive Performance Index (higher = worse cognition). OL, oligodendrocytes.

We therefore queried per-cluster expression for each of the 38 cilia structural genes against the Allen Brain Cell Atlas whole-mouse-brain taxonomy (release 20241130, subclass level), recording both mean expression and detection rate and weighting each cluster by its estimated abundance in hippocampal tissue. Marker controls behaved as expected: Foxj1 and Tmem212 were detected in 99.4% and 99.7% of ependymal cells versus 3.3% and 0.2% of CA1 glutamatergic neurons, whereas Fibcd1 and Camk2a were detected in 97.2% and 100% of CA1 neurons. Of the 38 module genes, 27 were ependymal-restricted, with a median detection rate of 90.2% in ependymal cells against 1.1% in CA1 glutamatergic neurons and a median 366-fold expression specificity. Ependymal-exclusive control genes established an ambient-RNA floor of 6.1% detection in CA1, which only Ak7, Pih1d2 and Spag17 marginally exceeded. Three genes were genuinely co-expressed in CA1 glutamatergic neurons, and did so across all six CA1-ProS supertypes rather than within a single subcluster: Dnah9 (CA1 85.6%, ependymal 98.2%), Dnai4 (71.6% and 95.5%) and Kif9 (30.6% and 96.5%). Two additional genes, Cfap65 and Enkur, were detected in subicular and presubicular rather than CA1 neurons. Ccp110, Fbrsl1 and Lrrc39 were evaluated separately, because Ccp110 is a cell-cycle-regulated centriolar gene expressed by any dividing cell and the remaining two have no established ciliary function.

Because ependymal cells are sparse and are preferentially lost during tissue dissociation, abundance weighting was based on spatial rather than dissociation-derived cell counts. Ependymal cells represented 0.160% of hippocampal cells by spatial measurement but only 0.015% of cells recovered in dissociation-based libraries, a 10.7-fold under-recovery. Under spatial weighting, ependymal cells were the top predicted contributor for 26 of the 38 genes and accounted for a median 49.3% of the predicted bulk signal. The cilia structural module therefore derives predominantly from ependymal cells rather than from hippocampal neurons.

### The cilia module score is independent of ependymal tissue content

Because ependymal cells line the lateral ventricle immediately adjacent to the dorsal hippocampus, the quantity of adherent ventricular ependyma captured during dissection could in principle vary between animals and generate a coordinated block of co-regulated transcripts. We therefore tested whether the association between the module score and cognitive performance was attributable to ependymal tissue content rather than to ependymal ciliary gene expression. Marker panels indexing ependymal ciliary expression (Foxj1, Drc1, Rsph4a, Rsph1, Tmem212, Dnah5, Hydin, Ccdc40), ependymal cell identity (Vim, Hdc, Rarres2, Clic6) and choroid plexus content (Prlr, Ttr, Kcnj13, Aqp1, Otx2, Folr1) were computed as the mean of gene-wise z-scores across the 21 animals. Ccdc146, Spag17 and Dnah9 were excluded from all panels because they are module members. Spag6 and Ccdc39 were excluded from the ciliary panel because atlas attribution showed that neither is ependymal-restricted in mouse (Spag6 detection 0.6% in ependymal cells versus 15.3% in CA1; Ccdc39 57.8% in CA1).

The module score was unrelated to Foxj1 expression, the most direct single marker of ependymal content (rho = −0.016, p = 0.95), and only weakly related to a composite ependymal abundance score whose confidence interval included zero (rho = 0.434, p = 0.050). Adding each panel as a covariate alongside age left the partial correlation between module score and CPI essentially unchanged: rho = −0.811 for the ependymal ciliary panel, −0.773 for ependymal identity, −0.787 for ependymal abundance, −0.810 for Foxj1 alone and −0.819 for choroid plexus content, against rho = −0.810 controlling for age alone. All bootstrap confidence intervals excluded zero and all permutation p values were below 0.001. Variation in the amount of ependymal or choroid plexus tissue sampled therefore does not account for the association. Taken together with the cell-type attribution, these data indicate that the module indexes expression of a motile ciliary program within a numerically stable ependymal population rather than the abundance of the cells that carry it.

Notably, most individual ependymal-restricted markers showed no association with cognitive performance after age adjustment (Foxj1 rho = −0.066, Dnah5 −0.048, Drc1 −0.086, Rsph4a - 0.140, Ccdc40 −0.168, Rsph1 −0.251, Hydin −0.259), whereas Tmem212 did (rho = −0.682, p = 0.0009). Tmem212 was not among the 271 CPI-associated genes and was therefore not subject to the selection applied to the module, so this association is not inflated by gene selection; its mean expression nonetheless lies near the lower bound of the detected range, and it should be treated as a single-gene observation requiring independent confirmation. The cognition-associated signal is therefore not a uniform shift across the ependymal ciliary program but is concentrated within a subset of its constituent genes.

### Cell-type proportion estimates do not account for the cilia module score

To complement the marker gene approach and estimate the relative abundance of each hippocampal cell type across the 21 bulk-derived samples, we performed CIBERSORTx cell-type deconvolution. As an initial low-resolution confirmation^40^, deconvolution using a five-cell-type reference (astrocytes, neurons, oligodendrocytes, microglia, endothelial cells) confirmed that the estimated cellular composition was predominantly neuronal across all animals, regardless of cognitive performance. To further resolve neuronal subpopulations, we then applied a higher-resolution 14-cell-type deconvolution using a single-cell RNA-seq reference derived from the mouse hippocampus^41^ (**Supplemental Figures 7 and 8**). Deconvolution was successful for all 21 samples (mean correlation between mixture and estimated profile = 0.737, range 0.729-0.750; mean RMSE = 2.30).

Nine cell types yielded non-zero proportion estimates across all samples. The most abundant estimated populations were Mossy-Cyp26b1 neurons (mean absolute score 1.19), Granule-mature neurons (0.31), Granule-immature neurons (0.29), and GABA-Lhx6 interneurons (0.23). Astrocytes, oligodendrocytes, endothelial cells, microglia, and Neuroblast 1 populations were also detected at lower proportions. Cajal-Retzius cells, Neuroblast 2, and nIPC populations yielded zero estimates across all samples.

Partial Spearman correlations between cell-type proportions and CPI, controlling for age, identified Mossy-Cyp26b1 neurons as the only cell type with a significant association (partial rho = +0.45, nominal p = 0.044; Table 2). Higher estimated proportions of Mossy-Cyp26b1 neurons were associated with worse cognitive performance (higher CPI values), independent of age. However, neither Mossy-Cyp26b1 nor other cell type proportions reached significance after age correction, and no comparisons survived Benjamini-Hochberg multiple comparison correction (all padj > 0.39); the functional significance of this association therefore remains to be determined.

We note that the 14-cell-type reference used for deconvolution contains no ependymal population, so these estimates cannot evaluate ependymal proportion directly; that question is addressed by the marker-panel analysis above. Within the populations the reference does resolve, no individual cell-type proportion significantly correlated with the 38-gene cilia structural module score after controlling for age (all p > 0.18; **Table 2**), suggesting that the cilia structural transcriptomic signal is not driven by a proportional shift in any of the populations this reference resolves. Gene-level expression of the cilia structural genes was directionally consistent with respect to cognitive performance, in contrast to the functionally heterogeneous top-ranked gene set, which displayed a bidirectional pattern (**Supplemental Figure 9**).

## DISCUSSION

Cognitive aging is widely assumed to reflect a progressive biological decline, yet individuals exhibit substantial variability in functional outcomes. By integrating automated behavioral phenotyping with population-based modeling and transcriptomic analysis, this study demonstrates that cognitive decline is not an inevitable consequence of chronological aging but instead reflects disruption of specific molecular programs that diverge across individuals over time. Importantly, transcriptional signatures associated with cognitive trajectory were largely separable from age-dependent gene-expression changes and were strongly enriched for ciliary structural pathways, implicating ciliated cells at the ventricular surface as candidate biological determinants of cognitive heterogeneity during aging.

Human studies have established that cognitive aging is highly heterogeneous^42–44^, but translating this concept to preclinical models has been limited by methodological variability and inconsistent behavioral metrics^22,45^. Using automated, high-resolution behavioral phenotyping, we show that substantial variability in performance emerges early in life and progressively diverges with age, even in genetically uniform mice. This early and progressive divergence, occurring in genetically uniform mice under standardized conditions, indicates that the sources of cognitive heterogeneity are present from early life, well before traditional aging-related decline becomes apparent, though whether these sources reflect even earlier developmental origins^14^ remains an open question for future investigation. Reversal-learning measures were particularly robust, combining sensitivity to age and sex with minimal susceptibility to procedural variability. Because reversal learning engages executive processes dependent on hippocampal as well as prefrontal circuitry, these measures likely provide translationally relevant endpoints that are also affected in human aging and neurodegenerative disease^42–48^. Automated behavioral platforms, therefore, offer a means to capture individual cognitive trajectories with greater precision than traditional cognitive testing protocols.

Combining these behavioral trajectories with hippocampal transcriptomic data revealed that cognitive decline is associated with a coherent and reproducible gene-expression program, largely separable from chronological aging. Cognition-linked genes were fewer, more structured, and more stable across subjects than the broad transcriptional changes associated with age, indicating that cognitive impairment represents a specific molecular phenotype rather than a generalized consequence of age-related transcriptional drift. Stability-based prioritization using the CSI metric developed here reinforced this distinction, showing that cognition-associated genes are consistently reproducible across animals rather than driven by outlier effects or cohort-specific variability.

Ciliary structural and motility genes, cytoskeletal organization, and microtubule-based transport emerged as a central biological themes among genes tracking cognitive trajectory. Cilia of both the motile and non-motile classes contribute to central nervous system function, and much of the existing literature concerns the non-motile primary cilium, which acts as a specialized signaling organelle regulating receptor trafficking, intracellular signaling, synaptic plasticity, metabolic sensing, and autophagy within the nervous system^49–62^. Disrupting ciliary structure or trafficking impairs dendritic development, synaptic maintenance, and hippocampal-dependent learning, underscoring a critical role for cilia in cognitive function. Ciliary signaling has also been linked to neurodegenerative and neuropsychiatric disorders^53,57,59,63^. A well-characterized example from that literature is systemic endocrine signaling through osteocalcin and its receptor GPR158, both localized to neuronal primary cilia, which links peripheral physiology to cognitive regulation^53,60,64^.

The genes identified here, however, encode components of the motile axoneme, including outer dynein arms, radial spokes and microtubule inner proteins, structures that have no counterpart in the non-motile primary cilia of neurons. Together with their attribution to ependymal cells, this directs attention to the motile ciliated epithelium lining of the lateral ventricle rather than to neuronal ciliary signaling. The mechanisms by which ependymal ciliary gene expression might influence cognitive performance were not examined here and remain to be established.

Despite these advances, the role of ciliary biology in age-related cognitive heterogeneity has remained largely unexplored. Here, cilia-associated transcriptional programs were selectively enriched among genes tracking cognitive trajectory and were largely absent from age-associated gene sets, indicating that differences in ciliary gene expression are not a general feature of chronological aging but instead distinguish cognitively impaired individuals throughout the lifespan. No measure of ciliary structure or function was obtained here, so these transcriptional differences should not be equated with ciliary dysfunction. These results are consistent with a model in which variability in the ependymal motile ciliary program is associated with the divergence in cognitive outcomes, with early differences potentially shaping trajectories that manifest as functional decline later in life.

Secondary enrichment of immune and inflammatory pathways, as reported in our study, is compatible with inflammatory signaling acting upon the ciliary program rather than arising as a consequence of it. The present cross-sectional design cannot establish the direction of this relationship, and either ordering remains consistent with emerging models of reciprocal interaction between tissue dysfunction and neuroimmune activity^65–69^. These pathways may also provide a molecular basis for recent findings that hospital-treated infections, including those that do not enter the central nervous system, confer increased dementia risk over subsequent decades^70^. The co-occurrence of mitochondrial and metabolic pathways within these shared transcriptional categories further suggests that systemic aging processes interact with ciliary signaling networks to modulate vulnerability^71^.

The Cognitive Stability Index (CSI) developed here adds an additional dimension to our approach by prioritizing genes whose association with cognitive performance is reproducible across subjects, rather than driven by outlier effects or a small number of aberrant animals. Because the CSI selects on directional consistency, the uniform direction of the retained cilia structural genes is, in part, a property of the selection criterion rather than an independent observation. What selection does not impose is the composition of the retained set: 36 of the 38 genes are resilience-associated, with higher expression tracking preserved cognition, whereas the broader CPI|Age set from which they were drawn contains both resilience- and impairment-associated genes spanning immune and metabolic functions. By integrating effect size with reproducibility, the CSI metric operationalizes molecular resilience as the maintenance of coordinated cellular programs, rather than simply the absence of pathological change, and identifies a focused set of cilia structural genes, including Ak7, Cimip2c, Nme9 and Lrrc78, as the highest-confidence correlates of sustained cognitive function within this cohort. Daw1 carries the largest raw partial correlation of the module (|rho| = 0.79) but ranks lower by CSI because its leave-one-out estimates are the most variable, illustrating the distinction the index is designed to draw.

Together, the convergence of ciliary, immune, and mitochondrial pathways suggests that motile ciliated cells at the cerebrospinal fluid interface may act as integrative hubs linking systemic physiology to cognitive outcomes^53,54,59,72–75^. Ciliary function depends on microtubule motor activity and is responsive to metabolic and inflammatory conditions, and may therefore be particularly sensitive to age-related physiological stress. In this model, intrinsic variation in ciliary gene expression accompanies differences in baseline cognitive trajectory, while age-related metabolic and immune challenges modulate progression toward impairment. The model offers a testable framework for the progressive divergence in cognitive performance with age in the absence of overt neurodegeneration (**Figure 4**). The direction of this relationship was not addressed here, and an alternative account in which inflammatory or metabolic state independently influences both ependymal ciliary transcription and cognitive performance remains equally consistent with the present data.

**Figure 4.**
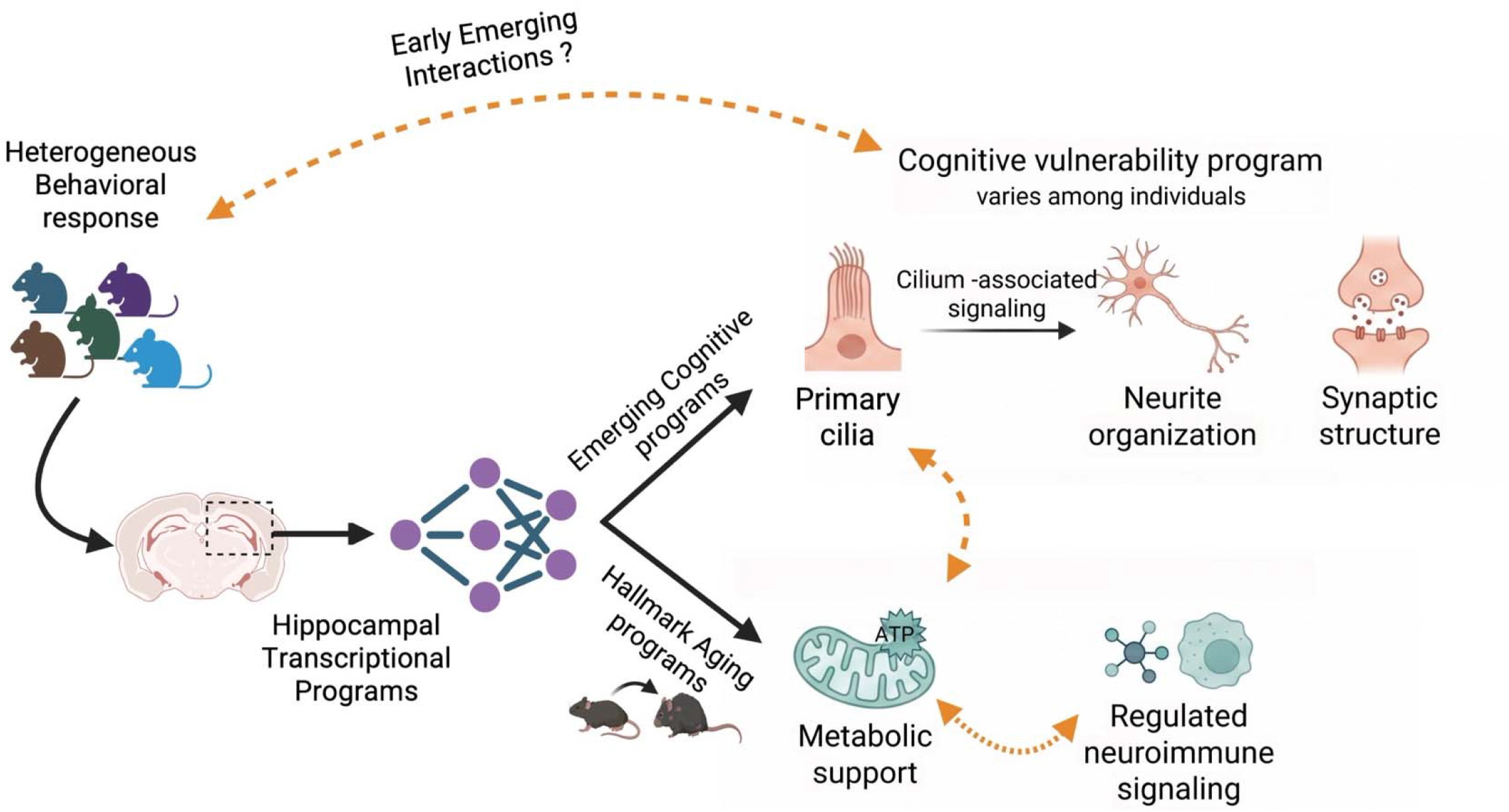
Conceptual model of age-independent and age-associated hippocampal transcriptional programs underlying individual differences in cognitive performance. Heterogeneous behavioral responses observed across isogenic C57BL/6 mice reflect underlying individual differences in hippocampal transcriptional organization. Bulk RNA-sequencing of hippocampal tissue from animals identified two distinct program classes. Emerging cognitive programs, defined by partial correlations with the Cognitive Performance Index independent of chronological age, converge on ciliary structural biology, with associated enrichment of neurite organization and synaptic structure pathways. Hallmark aging programs track age-dependent transcriptional shifts and are associated with metabolic support functions and regulated neuroimmune signaling. The dashed orange arrows indicate a hypothesized early-emerging "cognitive vulnerability program" that may establish individual differences in cognitive trajectories prior to the onset of detectable age-related decline, consistent with the Developmental Origins of Disease concept. Together, these programs suggest that cognitive heterogeneity in aging reflects the interaction of pre-existing molecular set points with superimposed aging processes. Cell-type attribution indicates that the ciliary component of this program derives predominantly from ependymal cells; the cilium shown is schematic and is not intended to specify a cell type. Figure created using BioRende

These findings carry important therapeutic implications. Interventions targeting broad aging processes may have limited impact on cognitive outcomes if they do not address the specific molecular pathways underlying cognitive impairment. Pathways centered on ciliary structure and intracellular transport, including the cilia structural genes identified through the CSI analysis (Daw1, Ccp110, Ak7, Dnai4, Cimip2c), represent more focused candidate targets for preserving cognitive function. Whether modulating these systems alters cognitive outcomes remains to be tested.

Beyond this specific finding, the analytical strategy developed here is broadly applicable to conditions characterized by heterogeneous cognitive outcomes. Cognitive impairment in neurodegenerative disease is not determined solely by pathological burden but instead reflects substantial individual variability consistent with cognitive reserve, cognitive resilience, and dissociation between clinical and pathological presentation. By deriving cognitive trajectories directly from behavioral performance and prioritizing molecular correlates based on stability, the framework presented here offers a generalizable approach for dissociating cognitive decline from aging or disease processes and for identifying pathways that selectively track vulnerability or resilience.

The cilia-cognition axis may represent one example of a broader phenomenon: the emergence of individualized, progressively diverging trajectories during aging. Because the behavioral cohort consisted of genetically identical C57BL/6 mice maintained under standardized conditions, the early appearance and progressive divergence of these trajectories cannot be attributed to genetic variation and likely reflect other sources of individual variation. Other domains of age-related decline, including those linked to more classical drivers such as oxidative stress, DNA damage and repair, and mitochondrial dysfunction, may follow similarly individualized trajectories that diverge before overt impairment is detectable, even where the underlying molecular pathways differ from those identified here for cognition.

The etiology of this early-emerging heterogeneity remains unknown. One possibility is that prenatal, perinatal, and early postnatal experiences, including maternal care^76–78^, litter size^16^, and early metabolic or physiological exposures^79,80^, establish lasting differences in neural and systemic set points that subsequently shape trajectories across the lifespan and contribute to longevity^81^. Under this view, the cilia-associated transcriptional differences identified in adulthood may represent a downstream readout of earlier developmental divergence rather than a process that originates *de novo* during aging. Longitudinal studies that track early-life conditions alongside behavioral and molecular trajectories will be required to test this possibility and could ultimately identify early-life windows for intervention.

Several limitations should be considered. The behavioral testing procedure required that we exclude animals that were not capable of performing the acquisition task to a 70% criteria. This resulted in the exclusion of animals that were blind or exhibit profound learning deficits. Although only 12% of animals were excluded, this likely represents another subgroup of animals with poor learning metrics and perhaps a unique cognitive trajectory not identified in the current results. Additionally, transcriptomic analyses were performed using bulk hippocampal tissue from a limited number of animals. Cellular origin was therefore inferred by attribution against an external reference rather than measured directly, and confirmation by in situ hybridization or immunohistochemistry distinguishing the CA1 pyramidal layer from the ventricular surface is required. A separate consideration is that the 38 module genes were selected on the basis of their association with cognitive performance in this same cohort, so the magnitude of the module-level partial correlation is subject to selection and should not be read as an unbiased effect-size estimate; calibration against a null set generated by rerunning the full selection pipeline on permuted cognitive performance values indicates that the association survives this correction. Reapplying the candidate-selection criteria within each of 2,000 permutations of the cognitive performance index and drawing gene sets of equal size from the resulting candidate pools yielded a median module-level partial rho of 0.45, with the observed value of −0.81 exceeded in 50 of 2,000 permutations (empirical p = 0.026); modelling the module as a co-expressed block within the candidate pool gave a median of 0.52 and an empirical p of 0.012 (Supplementary Figure 11). Roughly half of the observed magnitude is therefore attributable to the selection procedure, and the nominal p value substantially overstates the strength of the association. A separate control excluded transcript abundance as an explanation: the 38 module genes lie in the lowest decile of expression, but 2,000 abundance-matched gene sets drawn from genes not subject to selection gave a median partial rho of 0.17 and did not reach the observed value in any permutation (empirical p = 0.0005), and the magnitude of the gene-level partial correlation was unrelated to mean expression across the transcriptome (rho = - 0.02). Consistent with this, most individual ependymal ciliary genes that were not subject to selection show no association with cognitive performance. Finally, no measure of ciliary beating, cerebrospinal fluid flow or periventricular clearance was obtained here, so the functional consequence of the transcriptional differences remains untested, and establishing causal relationships between ciliary integrity and cognitive outcomes will require direct experimental validation.

Despite these limitations, our study is the first to utilize a large database of standardized behavioral testing in a mouse model that is robust across studies and timelines. The strength of this study is based on the cognitive performance index metric coupled with the permutation-validated transcriptional signature identified here, enriched for ciliary structure and motility, neurite organization, and synaptic architecture, with an empirical FDR of 5.4% and confirmed by leave-one-out stability analysis. These analyses provide a reproducible and empirically grounded foundation for mechanistic hypothesis testing. Together with the behavioral trajectory framework, these findings support a conceptual shift in how cognitive aging is understood: rather than an inevitable consequence of chronological age, cognitive trajectories appear to be shaped by variability in intrinsic molecular set points, with expression of ciliary structural pathways emerging as a candidate correlate of resilience. This integrated approach, linking trajectory-based behavioral phenotyping to stability-prioritized molecular correlates, offers a generalizable strategy for dissecting heterogeneous outcomes in aging, such as cognitive decline, and for identifying targets for interventions aimed at preserving function across the lifespan.

## METHODS

### Animals and behavioral testing

All procedures were approved by the Institutional Animal Care and Use Committee at the University of Oklahoma Health Sciences Center. Male and female C57BL/6N mice were obtained from the National Institute on Aging colony (Charles River Laboratories) and maintained under a 12-h light/12-h dark cycle at 21°C with *ad libitum* access to irradiated chow (5053 PicoLab, Purina Mills) and reverse-osmosis filtered water.

Behavioral data were derived from a longitudinal database comprising >3,400 animals aged 3– 30 months, including 1,240 animals initially characterized as controls. To minimize confounding, only vehicle-treated control animals were included, and animals undergoing invasive procedures were excluded. All studies were conducted using standardized housing and automated behavioral protocols between 2020 and 2024, enabling assessment of reproducibility across cohorts.

Cognitive testing was performed using an automated home-cage platform (PhenoTyper with CognitionWall; Noldus)^29–32,82–84^. Following acclimation, mice were trained to obtain a food reward by entering the left portal under a fixed-ratio 5 schedule (Acquisition phase). After 48 h, the rewarded location was switched to the right port (Reversal phase), requiring extinction of the learned response and acquisition of a new contingency. Animal position was recorded continuously for 89 h at 0.15-s intervals using infrared video tracking and processed with EthoVision XT14.^84,85^

Performance metrics were calculated as the percentage of correct responses over the trailing 30-trial window^31^. The number of entries and elapsed time required to reach performance thresholds (70%, 80%, and 90% correct) were quantified for both Acquisition and Reversal phases. Animals failing to reach ≥70% correct responses during initial discrimination were excluded to control for non-cognitive deficits. Based on these criteria, 906 animals were included in subsequent analyses (**Figure 1a**). Age groups (3–4, 5–6, 7–10, 20–22, 23–25, and 26–30 months) were defined to capture major lifespan transitions^82,86^ while maintaining adequate sample sizes.

### Analytical workflow

The analytical framework integrated behavioral phenotyping and transcriptomic analyses in sequential stages. Behavioral variables were normalized and evaluated using linear mixed-effects models to identify stable metrics across time. Dimensionality reduction and nonlinear trajectory modeling were applied to derive a continuous measure of cognitive performance independent of age. Gene expression analyses used partial correlation approaches to isolate cognition-dependent from age-dependent transcriptional programs. Stability-based metrics were then applied to prioritize robust gene–behavior associations.

### Behavioral data preprocessing and linear mixed-effects modeling

Behavioral variables exhibited strong positive skewness. Values exceeding the 95th percentile (∼2,000 entries or 40 h) were right-censored, and truncated distributions were fitted using Weibull, log-logistic, or gamma models. The best-fitting model (maximum likelihood) was used to impute censored values by sampling from the upper tail. Sensitivity analyses using alternative censoring thresholds (88%, 92%, and 97%) produced comparable results.

Because variables were non-normally distributed, transformation strategies were evaluated.^87^ The R-L/E metric was not adequately normalized by log transformation and was therefore rank-transformed and z-scored. Correlation matrices were used to identify collinear variables (r>0.75), which were excluded to prevent redundancy.

Linear mixed-effects models were used to quantify sources of variance. Random effects included Year and Run Date nested within Year, which provided the best model fit based on BIC. Fixed effects included Age, Sex, and Age × Sex interaction. Alternative random-effects structures (Year + Investigator/Project, Year + Mouse cohort) yielded comparable results. The test for zero variance for the random effect Year was based on the 50:50 mixture of a point mass at 0 and χ2 distribution.^35,36^ Models were implemented using SAS version 9.4.^37^

### Derivation of the Cognitive Performance Index (CPI)

Cognitive performance was assessed using three reversal-learning metrics (Re80, Rh80, and R-L/E), selected based on sensitivity and reproducibility. Variables were rank-normalized to minimize distributional assumptions.^88^

Principal component analysis identified a dominant behavioral axis (PC1; 84% variance explained) representing a unidimensional summary of performance. To capture nonlinear individual differences, UMAP was applied to the normalized variables using the uwot package in R with Euclidean distance, spectral initialization, n_neighbors = 15, min_dist = 0.01, and seed 42.^89–91^ A principal curve was fitted to the UMAP embedding using the princurve package in R, and arc-length distance along the curve was used to define a continuous cognitive trajectory.^92–97^ This value was rank-normalized, scaled from 0 to 20, and defined as the Cognitive Performance Index (CPI), with higher values indicating poorer cognitive performance.

Gaussian mixture models (mclust package, R) were applied to CPI distributions to identify discrete cognitive states. Models with four to six components were evaluated, and a four-component solution was selected based on BIC values and interpretability.^98,99^

### Animal selection for transcriptomic analysis

To identify gene expression patterns associated with individual differences in cognitive performance, a subset of 21 male animals was selected for hippocampal RNA-seq from a previously published dataset^100^ using an extreme-group design. Animals were drawn from the same cohort of 906 animals included in the behavioral meta-analysis and selected to enrich for transcriptomic signal at the poles of the CPI distribution. Specifically, animals were drawn from four groups defined by the intersection of age and CPI: young animals (5–6 months) with low CPI (n = 5, mean CPI = 1.79 ± 0.35) and high CPI (n = 3, mean CPI = 10.79 ± 1.10), and aged animals (21–25 months) with low CPI (n = 6, mean CPI = 5.83 ± 2.35) and high CPI (n = 7, mean CPI = 15.94 ± 1.35). All 21 animals were male, eliminating sex as a confounding variable in gene-behavior analyses. Because selection was based on CPI extremes rather than random sampling, the variance in CPI within this subset is enriched relative to a random sample, which increases sensitivity to detect gene-behavior associations but means that the effect sizes and p-values should be treated as hypothesis-generating rather than hypothesis-testing.

### RNA sequencing and preprocessing

RNA libraries were prepared using the Swift RNA kit and sequenced on an Illumina NovaSeq 6000 platform (2×150 bp reads). Raw reads were processed using fastp (v0.23.4) for quality control and adapter trimming. Transcript abundance was quantified using Salmon (v1.10.0) in selective alignment mode against the Ensembl GRCm39 mouse reference transcriptome. Transcript-level estimates were aggregated to gene-level counts using the tximport package and normalized using the Variance Stabilizing Transformation (VST) within the DESeq2 (v1.44.0) framework. VST-normalized values were used for all downstream correlation and expression analyses.

### Disentangling age- and cognition-associated gene expression

Spearman correlations were initially computed between each gene’s expression and either CPI or age across the 21 animals to characterize the overall landscape of gene-behavior associations (|r| ≥ 0.30, p < 0.05). To separate independent contributions of cognitive performance and age, partial Spearman correlations were computed: CPI | Age estimated the association between gene expression and CPI while controlling for age, and Age | CPI estimated the association between gene expression and age while controlling for CPI.^94,98,99,101,102^ Genes were classified as cognition-specific (significant CPI | Age, non-significant Age | CPI), age-specific (significant Age | CPI, non-significant CPI | Age), or shared (significant for both) based on these criteria.

To estimate the empirical false discovery rate, permutation testing was performed by randomly shuffling CPI values across animals (n = 1,000 permutations) while holding gene expression data constant. Partial Spearman correlations were recomputed for all 16,343 genes in each permutation, and the number of genes exceeding |r| ≥ 0.44 was recorded. An empirical FDR was calculated as the ratio of the mean number of genes passing threshold in permuted data to the number observed in the true data (**Supplemental Figure 4**). This approach accounts for the correlation structure of the transcriptomic data and the specific sample size, providing a more appropriate false discovery estimate than parametric correction methods that assume independence across tests.

### Stability Analysis and Cognitive Stability Index (CSI)

To assess the robustness of gene–behavior associations, leave-one-out (LOO) analyses were performed for genes with large partial Spearman effect sizes. For each gene, correlations were recalculated after sequential exclusion of each animal.^103,104^ Variability was summarized using median, interquartile range (IQR), and sign consistency across LOO iterations. A Cognitive Stability Index (CSI) was defined as: CSI = |rCPI|Age| × sign consistency × (1 − normalized IQR), where IQR values were normalized to the observed range to constrain CSI between 0 and 1. CSI was used for ranking and prioritization of high-confidence gene candidates rather than hypothesis testing. Genes surviving both permutation FDR filtering and the LOO stability filter were carried forward as high-confidence candidates.

### Classification of resilience- and impairment-associated genes

Genes were classified based on the direction of their CPI | Age partial correlation. Genes with negative correlations (higher expression associated with lower CPI, i.e., better cognitive performance) were defined as resilience-associated. Genes with positive correlations (higher expression associated with higher CPI, i.e., greater impairment) were defined as impairment-associated.

### Derivation of the CPI|Age gene set

A CPI|Age gene set was defined as the subset of transcriptome-wide genes meeting the following criteria in the partial Spearman analysis: absolute partial correlation coefficient |r| ≥ 0.44 and p ≤ 0.05 for the CPI | Age association, with the threshold of |r| ≥ 0.44 corresponding to a two-tailed significance level of approximately p < 0.05 for n = 21. This threshold was chosen to balance sensitivity with the constraint of the sample size. Applying these criteria yielded 271 genes (the CPI|Age gene set) whose expression was significantly associated with cognitive performance independent of age.

The CPI|Age gene set is functionally heterogeneous by design; it was derived empirically from transcriptome-wide partial correlations without prior functional filtering. The 271 genes include cilia structural and motility genes (e.g., axonemal dynein subunits, cilia-associated proteins), immune and inflammatory mediators, mitochondrial and metabolic genes, and a small number of olfactory receptors and expressed pseudogenes whose hippocampal co-regulation with CPI likely reflects statistical co-expression rather than direct mechanistic involvement. This heterogeneity is explicitly retained in the gene set and addressed analytically by examining gene-level correlations and cell-type attribution rather than treating the set as a functionally uniform module.

### Cilia gene module score

To specifically examine the relationship between cilia structural gene expression and cognitive performance, a cilia gene module score was computed using the 38 genes within the CPI|Age set with established roles in cilia assembly, intraflagellar transport, or axonemal structure and motility (*Ak7, Armc3, Ccdc113, Ccdc120, Ccdc146, Ccdc169, Ccdc170, Ccdc177, Ccdc180, Ccp110, Cfap52, Cfap65, Cfap77, Cimip2c, Cimip5, Daw1, Dnaaf1, Dnah11, Dnah3, Dnah6, Dnah9, Dnai4, Dynlt5, Enkur, Fbrsl1, Gas2l2, Iqca1, Kif9, Lrrc34, Lrrc39, Lrrc78, Morn5, Nme9, Odad2, Pih1d2, Spag16, Spag17, Ttc29*). The module score was defined as the mean VST-normalized expression value across these 38 genes for each animal. Partial Spearman correlations between the cilia module score and CPI, controlling for age, were computed using the ppcor package in R. Animals were stratified into cognitively impaired (high CPI) and cognitively resilient (low CPI) groups using a median split, and group differences in module score were assessed using a Welch two-sample t-test.

### Functional enrichment analysis

Functional enrichment was performed in R using the clusterProfiler (v4.12.3) and msigdbr (v25.1.1) packages. Gene Set Enrichment Analysis (GSEA) was conducted on the full transcriptome (16,343 genes) ranked by partial correlation coefficients (ρ) from the CPI | Age analysis, with results visualized via dotplots and enrichment plots. To illustrate the distinct transcriptional effects of aging and cognitive performance, a comparative GO analysis was performed across three models: Descriptive AGE, Inferential AGE | CPI (age adjusted for cognition), and CPI | AGE (cognition adjusted for age). Over-representation analysis (ORA) was performed on top gene subsets using a custom background universe of 16,343 genes. For all analyses, significance was defined by a Benjamini-Hochberg adjusted p-value < 0.05.

### Heatmap visualizations

Gene expression heatmaps were generated using the ComplexHeatmap (v2.20.0) R package. VST-normalized counts were Z-score standardized prior to visualization to ensure comparability across genes with different expression magnitudes. Samples were ordered by CPI and genes were ranked by their CPI | Age partial correlation coefficient (ρ). Separate heatmaps were generated for cognition-specific, age-specific, and shared gene sets to highlight their distinct expression trajectories.

### Cell-type deconvolution by CIBERSORTx

Cell-type deconvolution of the hippocampal RNA-seq data was performed using CIBERSORTx^105^; cibersortx.stanford.edu). A high-resolution 14-cell-type deconvolution was used to resolve distinct hippocampal neuronal and glial subpopulations and served as the primary deconvolution analysis. A single-cell RNA-seq library was obtained from the mouse hippocampus dataset of Zeisel et al^41^, GEO accession GSE95315, comprising 5,454 cells across 22 annotated cell types. Cell types with fewer than 66 cells were excluded, and the remaining types were subsampled to 66 cells each, yielding a balanced reference of 14 cell types (924 cells total). The signature matrix was constructed via the CIBERSORTx web portal in single-cell reference mode with the following settings: replicates = 5; sampling fraction = 0.5; maximum condition number (kappa) = 999; q-value threshold = 0.01; quantile normalization disabled. The resulting signature matrix comprised 14 cell-type columns across 14,545 genes: Astrocytes, Cajal-Retzius, Endothelial, GABA-Lhx6, Granule-immature, Granule-mature, Microglia, Mossy-Cyp26b1, Neuroblast_1, Neuroblast_2, nIPC, OL, OPC, and Radial_Glia-like.

Cell fraction imputation was performed in absolute mode with B-mode batch correction to account for platform differences between the single-cell reference (10x Chromium) and the bulk

RNA-seq data. Bulk expression values were back-transformed from VST-normalized space to linear space prior to submission as the mixture file. One hundred permutations were used for significance testing; all 21 samples returned statistically significant deconvolution solutions (p < 0.05). Estimated cell-type proportions were merged with animal-level metadata, including age and CPI. Partial Spearman correlations between each cell-type proportion and both the cilia module score and CPI were computed, controlling for age, using the ppcor package in R, with Benjamini-Hochberg correction applied for multiple comparisons.

### Cell-type attribution of CPI | Age genes

To characterize the cell-type context of CPI|Age gene expression, attribution was performed using two complementary approaches. First, the Allen Brain Cell Atlas (ABCA) whole-mouse-brain transcriptomic taxonomy^106^ (Allen Institute for Brain Science, 2023; portal.brain-map.org) was queried for direct single-cell-based subclass assignments. Cell type cluster annotations were downloaded from the ABC Atlas AWS public dataset (version 20231215). Hippocampal-associated clusters were identified by filtering on CCF anatomical acronyms (HIP, CA1, CA2, CA3, DG, HPF), and each gene in the CPI|Age set was matched against cluster marker fields using exact word-boundary matching. This procedure yielded direct subclass labels for 18 of the 271 genes across 13 cell subclasses, with binary flags for neuronal identity and hippocampal specificity.

For the remaining 253 genes lacking direct ABCA assignments, cell-type attribution was derived empirically by correlating each gene’s expression across the 21 animals against each of the 14 CIBERSORTx-estimated cell-type proportions using Spearman rank correlation. Each gene was assigned to the cell type whose proportion most strongly co-varied with its expression (highest absolute Spearman ρ). The strength of each attribution was retained as a quality metric (e.g. best_ρ); genes with best_ρ < 0.3 were considered weakly attributed and interpreted with caution. This approach is most informative for cell-type-enriched transcripts and represents a probabilistic rather than definitive assignment, particularly for broadly expressed genes.

### Gene-level CPI correlation and cell-type enrichment analysis

Spearman rank correlations were computed between the VST-normalized expression of each of the 271 CPI|Age genes and animal CPI scores across all 21 animals using the cor.test function in R (v4.4.1; method = ’spearman’, exact = FALSE). This analysis was performed without age correction to characterize the raw expression-CPI relationship for each gene, complementing the partial correlations used to derive the gene set. Multiple testing correction was applied using the Benjamini-Hochberg FDR procedure. Genes were classified as FDR-significant at p-adjusted < 0.05. Genes with ρ > 0.3 were considered impairment-associated and those with ρ < −0.3 were considered resilience-associated, thresholds corresponding to a moderate-to-strong effect size for this sample size.

To assess whether differences in estimated cell-type composition across animals could account for gene-CPI associations, Spearman correlations between each of the 14 CIBERSORTx cell-type proportions and CPI were computed as a specificity control, with Benjamini-Hochberg corrections applied. Genes were then grouped by derived cell-type attribution, and summary statistics were computed per group: number of genes, mean and median Spearman ρ, number of genes with ρ > 0.3 or ρ < −0.3, and number nominally significant at p < 0.05. Summaries were computed separately for ABCA-derived and CIBERSORTx-derived attributions.

### Statistical analysis

All statistical analyses were performed in R (v4.4.1) or SAS version 9.4. Tests were two-sided with significance defined as p < 0.05 unless otherwise noted. Spearman rank correlations were used throughout for gene-behavior and cell-type analyses, given non-normal distributions and the modest sample size (n = 21 for transcriptomic analyses). Age was treated as a continuous covariate in all partial correlation analyses. Multiple testing correction used the Benjamini-Hochberg method where specified. Linear mixed-model assumptions were evaluated using residual diagnostics. No formal power calculations were performed prior to study initiation. Investigators were not blinded to group assignments, since analyses were conducted using automated pipelines with predefined inclusion criteria. Figures were generated using Prism (v11.0.2) or ggplot2 (v4.0.2) and patchwork packages in R.

### Data and code availability

Behavioral data and analysis scripts are available from the corresponding author upon reasonable request. RNA-seq raw data files from this study can be accessed through the NCBI Sequence Read Archive under the BioProject: PRJNA1483660. R analysis code for all gene-level correlations and analysis is available upon request.

## ACKNOWLEDGEMENTS

This work was supported by the National Institute on Aging, the National Institute of General Medical Sciences, and National Institute of General Medical Sciences and National Institute of Aging of the National Institutes of Health under award numbers 2P20GM125528 (WES) and 5T32AG052363 (WES); NIA R01AG085398 (SL) and Hevolution/AFAR New Investigator Award (SL). The content is solely the responsibility of the authors and does not necessarily represent the official views of the NIH. Additional support was provided by a Presbyterian Health Foundation Grant. We gratefully acknowledge the contributions of Kylie Williams and Melissa K. Day in the preparation of this manuscript.

## AUTHOR CONTRIBUTIONS

Drs. Logan and Sonntag designed the behavioral phenotyping studies, conducted the behavioral analyses, and wrote the initial draft of the manuscript. Dr. Sonntag also performed initial statistical analyses confirming stability of the behavioral measures across the 5-year period. Dr. Ding served as primary statistician, analyzing and/or validating all statistical assessments. Dr. Baier performed initial transformation of behavioral data into UMAP and Gaussian distributions and contributed to manuscript writing. Drs. Unnikrishnan and Diaz-Garcia contributed to overall study design and data analysis. Dr. Drevets contributed to manuscript writing and overall direction. Drs. Kurup, Jung, and Oh provided bioinformatics expertise and analyzed gene expression data.

## REFERENCES

1 Rapp, P. R. & Amaral, D. G. Individual differences in the cognitive and neurobiological consequences of normal aging. Trends Neurosci 15, 340–345 (1992). 10.1016/0166-2236(92)90051-9

2 Mungas, D. et al. Heterogeneity of cognitive trajectories in diverse older persons. Psychol Aging 25, 606–619 (2010). 10.1037/a0019502

3 Gray, D. T., Smith, A. C., Burke, S. N., Gazzaley, A. & Barnes, C. A. Attentional updating and monitoring and affective shifting are impacted independently by aging in macaque monkeys. Behav Brain Res 322, 329–338 (2017). 10.1016/j.bbr.2016.06.056

4 McQuail, J. A. et al. Cognitive Reserve in Model Systems for Mechanistic Discovery: The Importance of Longitudinal Studies. Front Aging Neurosci 12, 607685 (2020). 10.3389/fnagi.2020.607685

5 Barzilai, N., Atzmon, G., Derby, C. A., Bauman, J. M. & Lipton, R. B. A genotype of exceptional longevity is associated with preservation of cognitive function. Neurology 67, 2170–2175 (2006). 10.1212/01.wnl.0000249116.50854.65

6 Bergman, A., Atzmon, G., Ye, K., MacCarthy, T. & Barzilai, N. Buffering mechanisms in aging: a systems approach toward uncovering the genetic component of aging. PLoS Comput Biol 3, e170 (2007). 10.1371/journal.pcbi.0030170

7 Stern, Y. Cognitive reserve in ageing and Alzheimer’s disease. Lancet Neurol 11, 1006– 1012 (2012). 10.1016/S1474-4422(12)70191-6

8 Aarsland, D. et al. Mild cognitive impairment in Parkinson disease: a multicenter pooled analysis. Neurology 75, 1062–1069 (2010). 10.1212/WNL.0b013e3181f39d0e

9 Bennett, D. A. et al. Neuropathology of older persons without cognitive impairment from two community-based studies. Neurology 66, 1837–1844 (2006). 10.1212/01.wnl.0000219668.47116.e6

10 Boyle, P. A., Yu, L., Wilson, R. S., Schneider, J. A. & Bennett, D. A. Relation of neuropathology with cognitive decline among older persons without dementia. Front Aging Neurosci 5, 50 (2013). 10.3389/fnagi.2013.00050

11 Snowdon, D. A. et al. Linguistic ability in early life and cognitive function and Alzheimer’s disease in late life. Findings from the Nun Study. JAMA 275, 528–532 (1996).

12. Kuchel, G. A., Hevener, A. L., Ruby, J. G., Sebastiani, P. & Kumar, V. Workshop Report-Heterogeneity and Successful Aging Part I: Heterogeneity in Aging-Challenges and Opportunities. J Gerontol A Biol Sci Med Sci 80 (2025). 10.1093/gerona/glaf023

13. Kumar, V., et al. Workshop Report-Heterogeneity and Successful Aging Part II: Approaches to Investigate Heterogeneity in Aging Research. J Gerontol A Biol Sci Med Sci 80 (2025). 10.1093/gerona/glaf021

14 Barker, D. J. The developmental origins of adult disease. Eur J Epidemiol 18, 733–736 (2003). 10.1023/a:1025388901248

15 Huang, Z., Jordan, J. D. & Zhang, Q. Early life adversity as a risk factor for cognitive impairment and Alzheimer’s disease. Transl Neurodegener 12, 25 (2023). 10.1186/s40035-023-00355-z

16 Krejcova, L. V. et al. Unraveling the Influence of Litter Size, Maternal Care, Exercise, and Aging on Neurobehavioral Plasticity and Dentate Gyrus Microglia Dynamics in Male Rats. Brain Sci 14 (2024). 10.3390/brainsci14050497

17 Kuhn, H. G., Skau, S. & Nyberg, J. A lifetime perspective on risk factors for cognitive decline with a special focus on early events. Cereb Circ Cogn Behav 6, 100217 (2024). 10.1016/j.cccb.2024.100217

18 Kunzi, M., Gheorghe, D. A., Gallacher, J. & Bauermeister, S. The impact of early adversity on later life health, lifestyle, and cognition. BMC Public Health 24, 3294 (2024). 10.1186/s12889-024-20768-3

19 Raikkonen, K. et al. Early life origins cognitive decline: findings in elderly men in the Helsinki Birth Cohort Study. PLoS One 8, e54707 (2013). 10.1371/journal.pone.0054707

20 Short, A. K. & Baram, T. Z. Early-life adversity and neurological disease: age-old questions and novel answers. Nat Rev Neurol 15, 657–669 (2019). 10.1038/s41582-019-0246-5

21 Shoji, H., Takao, K., Hattori, S. & Miyakawa, T. Age-related changes in behavior in C57BL/6J mice from young adulthood to middle age. Mol Brain 9, 11 (2016). 10.1186/s13041-016-0191-9

22 Sare, R. M., Lemons, A. & Smith, C. B. Behavior Testing in Rodents: Highlighting Potential Confounds Affecting Variability and Reproducibility. Brain Sci 11 (2021). 10.3390/brainsci11040522

23 van der Goot, M. H. et al. Incorporating inter-individual variability in experimental design improves the quality of results of animal experiments. PLoS One 16, e0255521 (2021). 10.1371/journal.pone.0255521

24 Begley, C. G. & Ellis, L. M. Drug development: Raise standards for preclinical cancer research. Nature 483, 531–533 (2012). 10.1038/483531a

25 Bernstein, B. J. et al. Sex differences in spontaneous behavior and cognition in mice using an automated behavior monitoring system. Physiol Behav 283, 114595 (2024). 10.1016/j.physbeh.2024.114595

26 Endo, T. et al. Automated test of behavioral flexibility in mice using a behavioral sequencing task in IntelliCage. Behav Brain Res 221, 172–181 (2011). 10.1016/j.bbr.2011.02.037

27 Grieco, F. et al. Measuring Behavior in the Home Cage: Study Design, Applications, Challenges, and Perspectives. Front Behav Neurosci 15, 735387 (2021). 10.3389/fnbeh.2021.735387

28 Ho, H. et al. A fully automated home cage for long-term continuous phenotyping of mouse cognition and behavior. Cell Rep Methods 3, 100532 (2023). 10.1016/j.crmeth.2023.100532

29 Loos, M. et al. Sheltering behavior and locomotor activity in 11 genetically diverse common inbred mouse strains using home-cage monitoring. PLoS One 9, e108563 (2014). 10.1371/journal.pone.0108563

30 Mingrone, A., Kaffman, A. & Kaffman, A. The Promise of Automated Home-Cage Monitoring in Improving Translational Utility of Psychiatric Research in Rodents. Front Neurosci 14, 618593 (2020). 10.3389/fnins.2020.618593

31 Remmelink, E., Smit, A. B., Verhage, M. & Loos, M. Measuring discrimination- and reversal learning in mouse models within 4 days and without prior food deprivation. Learn Mem 23, 660–667 (2016). 10.1101/lm.042085.116

32 Richter, S. H. Automated Home-Cage Testing as a Tool to Improve Reproducibility of Behavioral Research? Front Neurosci 14, 383 (2020). 10.3389/fnins.2020.00383

33 Ruby, J. G. et al. An Automated, Home-Cage, Video Monitoring-based Mouse Frailty Index Detects Age-associated Morbidity in C57BL/6 and Diversity Outbred Mice. J Gerontol A Biol Sci Med Sci 78, 762–770 (2023). 10.1093/gerona/glad035

34 Voikar, V. & Gaburro, S. Three Pillars of Automated Home-Cage Phenotyping of Mice: Novel Findings, Refinement, and Reproducibility Based on Literature and Experience. Front Behav Neurosci 14, 575434 (2020). 10.3389/fnbeh.2020.575434

35 Fitzmaurice, G. M., Laird, N. M., & Ware, J. H. . Applied longitudinal analysis. (John Wiley & Sons, 2012).

36 Laird, N. M. & Ware, J. H. Random-effects models for longitudinal data. Biometrics 38, 963–974 (1982).

37 Stroup, W. W., Milliken, G. A., Claassen, E. A., & Wolfinger, R. D. (SAS Institute, 2018).

38 Lopez-Otin, C., Blasco, M. A., Partridge, L., Serrano, M. & Kroemer, G. The hallmarks of aging. Cell 153, 1194–1217 (2013). 10.1016/j.cell.2013.05.039

39 Lopez-Otin, C., Blasco, M. A., Partridge, L., Serrano, M. & Kroemer, G. Hallmarks of aging: An expanding universe. Cell 186, 243–278 (2023). 10.1016/j.cell.2022.11.001

40 Zhang, Y. et al. An RNA-sequencing transcriptome and splicing database of glia, neurons, and vascular cells of the cerebral cortex. J Neurosci 34, 11929–11947 (2014). 10.1523/JNEUROSCI.1860-14.2014

41 Zeisel, A. et al. Molecular Architecture of the Mouse Nervous System. Cell 174, 999– 1014 e1022 (2018). 10.1016/j.cell.2018.06.021

42 Evans, D. A., Grodstein, F., Loewenstein, D., Kaye, J. & Weintraub, S. Reducing case ascertainment costs in U.S. population studies of Alzheimer’s disease, dementia, and cognitive impairment-Part 2. Alzheimers Dement 7, 110–123 (2011). 10.1016/j.jalz.2010.11.008

43 Harrison, T. M., Weintraub, S., Mesulam, M. M. & Rogalski, E. Superior memory and higher cortical volumes in unusually successful cognitive aging. J Int Neuropsychol Soc 18, 1081–1085 (2012). 10.1017/S1355617712000847

44 Weintraub, S., Gefen, T., Geula, C. & Mesulam, M. M. The first 25 years of the Northwestern University SuperAging Program. Alzheimers Dement 21, e70312 (2025). 10.1002/alz.70312

45 Brito, D. V. C. et al. Assessing cognitive decline in the aging brain: lessons from rodent and human studies. NPJ Aging 9, 23 (2023). 10.1038/s41514-023-00120-6

46 Bagley, J. R. et al. Behavioral phenotypes revealed during reversal learning are linked with novel genetic loci in diversity outbred mice. Addict Neurosci 4 (2022). 10.1016/j.addicn.2022.100045

47 Izquierdo, A., Brigman, J. L., Radke, A. K., Rudebeck, P. H. & Holmes, A. The neural basis of reversal learning: An updated perspective. Neuroscience 345, 12–26 (2017). 10.1016/j.neuroscience.2016.03.021

48 Weiler, J. A., Bellebaum, C. & Daum, I. Aging affects acquisition and reversal of reward-based associative learning. Learn Mem 15, 190–197 (2008). 10.1101/lm.890408

49 Amador-Arjona, A. et al. Primary cilia regulate proliferation of amplifying progenitors in adult hippocampus: implications for learning and memory. J Neurosci 31, 9933–9944 (2011). 10.1523/JNEUROSCI.1062-11.2011

50 Armato, U., Chakravarthy, B., Pacchiana, R. & Whitfield, J. F. Alzheimer’s disease: an update of the roles of receptors, astrocytes and primary cilia (review). Int J Mol Med 31, 3–10 (2013). 10.3892/ijmm.2012.1162

51 Berbari, N. F. et al. Hippocampal and cortical primary cilia are required for aversive memory in mice. PLoS One 9, e106576 (2014). 10.1371/journal.pone.0106576

52 Chakravarthy, B. et al. Reduction of the immunostainable length of the hippocampal dentate granule cells’ primary cilia in 3xAD-transgenic mice producing human Abeta(1-42) and tau. Biochem Biophys Res Commun 427, 218–222 (2012). 10.1016/j.bbrc.2012.09.056

53 Guemez-Gamboa, A., Coufal, N. G. & Gleeson, J. G. Primary cilia in the developing and mature brain. Neuron 82, 511–521 (2014). 10.1016/j.neuron.2014.04.024

54 Huang, A. S., Li, A. S. M. & Hor, C. H. H. Primary cilia in the mature brain: emerging roles in Alzheimer’s disease pathogenesis. Front Cell Dev Biol 13, 1650884 (2025). 10.3389/fcell.2025.1650884

55 Karunakaran, K. B., Chaparala, S., Lo, C. W. & Ganapathiraju, M. K. Cilia interactome with predicted protein-protein interactions reveals connections to Alzheimer’s disease, aging and other neuropsychiatric processes. Sci Rep 10, 15629 (2020). 10.1038/s41598-020-72024-4

56 Khrimian, L. et al. Gpr158 mediates osteocalcin’s regulation of cognition. J Exp Med 214, 2859–2873 (2017). 10.1084/jem.20171320

57 Kobayashi, Y. et al. Impairment of ciliary dynamics in an APP knock-in mouse model of Alzheimer’s disease. Biochem Biophys Res Commun 610, 85–91 (2022). 10.1016/j.bbrc.2022.04.050

58 Loukil, A., Barrington, C. & Goetz, S. C. A complex of distal appendage-associated kinases linked to human disease regulates ciliary trafficking and stability. Proc Natl Acad Sci U S A 118 (2021). 10.1073/pnas.2018740118

59 Loukil, A. et al. Identification of New Ciliary Signaling Pathways in the Brain and Insights into Neurological Disorders. J Neurosci 45 (2025). 10.1523/JNEUROSCI.0800-24.2025

60 Obri, A., Khrimian, L., Karsenty, G. & Oury, F. Osteocalcin in the brain: from embryonic development to age-related decline in cognition. Nat Rev Endocrinol 14, 174–182 (2018). 10.1038/nrendo.2017.181

61 Rhee, S., Kirschen, G. W., Gu, Y. & Ge, S. Depletion of primary cilia from mature dentate granule cells impairs hippocampus-dependent contextual memory. Sci Rep 6, 34370 (2016). 10.1038/srep34370

62 Rivagorda, M. et al. A primary cilia-autophagy axis in hippocampal neurons is essential to maintain cognitive resilience. Nat Aging 5, 450–467 (2025). 10.1038/s43587-024-00791-0

63 Youn, Y. H. & Han, Y. G. Primary Cilia in Brain Development and Diseases. Am J Pathol 188, 11–22 (2018). 10.1016/j.ajpath.2017.08.031

64 Blanchet, V., Oury, F. & Romeo-Guitart, D. Neuronal antenna senses signals from the bone to sustain cognition by boosting autophagy. Autophagy 21, 1850–1852 (2025). 10.1080/15548627.2025.2487038

65 Andersen, J. V. et al. Hippocampal disruptions of synaptic and astrocyte metabolism are primary events of early amyloid pathology in the 5xFAD mouse model of Alzheimer’s disease. Cell Death Dis 12, 954 (2021). 10.1038/s41419-021-04237-y

66 Cali, C., Cantando, I., Veloz Castillo, M. F., Gonzalez, L. & Bezzi, P. Metabolic Reprogramming of Astrocytes in Pathological Conditions: Implications for Neurodegenerative Diseases. Int J Mol Sci 25 (2024). 10.3390/ijms25168922

67 Garrahy, J. P. Astrocytic lipidopathy and bioenergetic failure in ApoE4-associated late-onset Alzheimer’s disease: A unifying hypothesis. J Alzheimers Dis 106, 890–902 (2025). 10.1177/13872877251350338

68 Kriska, J., Hermanova, Z., Knotek, T., Tureckova, J. & Anderova, M. On the Common Journey of Neural Cells through Ischemic Brain Injury and Alzheimer’s Disease. Int J Mol Sci 22 (2021). 10.3390/ijms22189689

69 Picon-Galindo, E., Latz, E. & Wachten, D. Primary cilia and their effects on immune cell functions and metabolism: a model. Trends Immunol 43, 366–378 (2022). 10.1016/j.it.2022.03.001

70 Lee, J. W. et al. Impact of multiple infections on risk of incident dementia according to subjective cognitive decline status: a nationwide population-based cohort study. Front Aging Neurosci 16, 1410185 (2024). 10.3389/fnagi.2024.1410185

71 Ignatenko, O. et al. Mitochondrial dysfunction compromises ciliary homeostasis in astrocytes. J Cell Biol 222 (2023). 10.1083/jcb.202203019

72 Finetti, F., Onnis, A. & Baldari, C. T. Regulation of vesicular traffic at the T cell immune synapse: lessons from the primary cilium. Traffic 16, 241–249 (2015). 10.1111/tra.12241

73 Ma, N. & Zhou, J. Functions of Endothelial Cilia in the Regulation of Vascular Barriers. Front Cell Dev Biol 8, 626 (2020). 10.3389/fcell.2020.00626

74 Nachury, M. V. & Mick, D. U. Establishing and regulating the composition of cilia for signal transduction. Nat Rev Mol Cell Biol 20, 389–405 (2019). 10.1038/s41580-019-0116-4

75 Paul, C. et al. Loss of primary cilia promotes inflammation and carcinogenesis. EMBO Rep 23, e55687 (2022). 10.15252/embr.202255687

76 Glynn, L. M., Howland, M. A. & Fox, M. Maternal programming: Application of a developmental psychopathology perspective. Dev Psychopathol 30, 905–919 (2018). 10.1017/S0954579418000524

77 Lauby, S. C. et al. Postnatal maternal care moderates the effects of prenatal bisphenol exposure on offspring neurodevelopmental, behavioral, and transcriptomic outcomes. bioRxiv (2023). 10.1101/2023.09.19.558481

78 Weaver, I. C. et al. Epigenetic programming by maternal behavior. Nat Neurosci 7, 847– 854 (2004). 10.1038/nn1276

79 Khambadkone, S. G., Cordner, Z. A. & Tamashiro, K. L. K. Maternal stressors and the developmental origins of neuropsychiatric risk. Front Neuroendocrinol 57, 100834 (2020). 10.1016/j.yfrne.2020.100834

80 Sanchez-Garrido, M. A., Garcia-Galiano, D. & Tena-Sempere, M. Early programming of reproductive health and fertility: novel neuroendocrine mechanisms and implications in reproductive medicine. Hum Reprod Update 28, 346–375 (2022). 10.1093/humupd/dmac005

81 Moore, E. et al. Maternal obesity decreases offspring lifespan. Mol Metab 105, 102333 (2026). 10.1016/j.molmet.2026.102333

82 Logan, S. et al. Simultaneous assessment of cognitive function, circadian rhythm, and spontaneous activity in aging mice. Geroscience 40, 123–137 (2018). 10.1007/s11357-018-0019-x

83 Logan, S. et al. Accelerated decline in cognition in a mouse model of increased oxidative stress. Geroscience 41, 591–607 (2019). 10.1007/s11357-019-00105-y

84 Noldus, L. P., Spink, A. J. & Tegelenbosch, R. A. EthoVision: a versatile video tracking system for automation of behavioral experiments. Behav Res Methods Instrum Comput 33, 398–414 (2001). 10.3758/bf03195394

85 Spink, A. J., Tegelenbosch, R. A., Buma, M. O. & Noldus, L. P. The EthoVision video tracking system--a tool for behavioral phenotyping of transgenic mice. Physiol Behav 73, 731–744 (2001). 10.1016/s0031-9384(01)00530-3

86 Ashpole, N. M. et al. IGF-1 has sexually dimorphic, pleiotropic, and time-dependent effects on healthspan, pathology, and lifespan. Geroscience 39, 129–145 (2017). 10.1007/s11357-017-9971-0

87 Wang, B., Sun, F. & Luan, Y. Comparison of the effectiveness of different normalization methods for metagenomic cross-study phenotype prediction under heterogeneity. Sci Rep 14, 7024 (2024). 10.1038/s41598-024-57670-2

88 Zuur, A. F., Ieno, E. N. & Elphick, C. S. A protocol for data exploration to avoid common statistical problems. Methods in Ecology and Evolution 1, 3–14 (2010). 10.1111/j.2041-210X.2009.00001.x

89 Becht, E. et al. Dimensionality reduction for visualizing single-cell data using UMAP. Nat Biotechnol (2018). 10.1038/nbt.4314

90 Li, Y. et al. Pursuing sources of heterogeneity in modeling clustered population. Biometrics 78, 716–729 (2022). 10.1111/biom.13434

91 Nguyen, T. T. T. & Chang, Y. L. Insights into the Heterogeneity of Cognitive Aging: A Comparative Analysis of Two Data-Driven Clustering Algorithms. J Gerontol B Psychol Sci Soc Sci 80 (2025). 10.1093/geronb/gbaf022

92 Campbell, K. R. & Yau, C. Probabilistic modeling of bifurcations in single-cell gene expression data using a Bayesian mixture of factor analyzers. Wellcome Open Res 2, 19 (2017). 10.12688/wellcomeopenres.11087.1

93 Campbell, K. R. & Yau, C. A descriptive marker gene approach to single-cell pseudotime inference. Bioinformatics 35, 28–35 (2019). 10.1093/bioinformatics/bty498

94 Liu, J., Yang, M., Zhao, W. & Zhou, X. CCPE: cell cycle pseudotime estimation for single cell RNA-seq data. Nucleic Acids Res 50, 704–716 (2022). 10.1093/nar/gkab1236

95 Liu, Z. et al. Reconstructing cell cycle pseudo time-series via single-cell transcriptome data. Nat Commun 8, 22 (2017). 10.1038/s41467-017-00039-z

96 Trapnell, C. et al. The dynamics and regulators of cell fate decisions are revealed by pseudotemporal ordering of single cells. Nat Biotechnol 32, 381–386 (2014). 10.1038/nbt.2859

97 von Ziegler, L., Sturman, O. & Bohacek, J. Big behavior: challenges and opportunities in a new era of deep behavior profiling. Neuropsychopharmacology 46, 33–44 (2021). 10.1038/s41386-020-0751-7

98 Kim, N., Heo, M., Fleysher, R., Branch, C. A. & Lipton, M. L. A gaussian mixture model approach for estimating and comparing the shapes of distributions of neuroimaging data: diffusion-measured aging effects in brain white matter. Front Public Health 2, 32 (2014). 10.3389/fpubh.2014.00032

99 Schafer, J. & Strimmer, K. An empirical Bayes approach to inferring large-scale gene association networks. Bioinformatics 21, 754–764 (2005). 10.1093/bioinformatics/bti062

100 Logan, S. et al. Cognitive heterogeneity reveals molecular signatures of age-related impairment. PNAS Nexus 2, pgad101 (2023). 10.1093/pnasnexus/pgad101

101 Slinker, B. K. & Glantz, S. A. Multiple regression for physiological data analysis: the problem of multicollinearity. Am J Physiol 249, R1–12 (1985). 10.1152/ajpregu.1985.249.1.R1

102 Vatcheva, K. P., Lee, M., McCormick, J. B. & Rahbar, M. H. Multicollinearity in Regression Analyses Conducted in Epidemiologic Studies. Epidemiology (Sunnyvale) 6 (2016). 10.4172/2161-1165.1000227

103 Davison AC, H. D. Bootstrap Methods and Their Application. (Cambridge University Press, 1997).

104 Efron B, T. R. An Introduction to the Bootstrap., (Chapman & Hall/CRC, 1993).

105 Newman, A. M. et al. Determining cell type abundance and expression from bulk tissues with digital cytometry. Nat Biotechnol 37, 773–782 (2019). 10.1038/s41587-019-0114-2

106 Yao, Z. et al. A high-resolution transcriptomic and spatial atlas of cell types in the whole mouse brain. Nature 624, 317–332 (2023). 10.1038/s41586-023-06812-z

