## supplemental files 1-11 for "COGNITIVE HETEROGENEITY AND AGING: AN AGE-INDEPENDENT CILIARY GENE PROGRAM TRACKS COGNITIVE RESILIENCE IN MICE"

**SUPPLEMENT TABLE 1**

**Table 1: Barriers to Assessing the Mechanisms of Cognitive Decline in Humans**

1. **Limited Access to Human Brain Tissue:** The ability to procure brain samples from well-defined subsets of individuals with demonstrated cognitive impairment has improved but remains a limitation in the field.
2. **Invasiveness and Ethical Barriers**: Direct examination of living human brain tissue (e.g., via biopsy) is highly invasive and ethically constrained. Most cellular-level studies rely on post-mortem tissue, which only provides a snapshot after extensive disease progression or end-stage pathological changes.
3. **Post-mortem Artefacts**: Changes that occur after death (e.g., protein degradation, tissue damage) can confound interpretation of underlying mechanisms.
4. **Complexity and Heterogeneity of the Human Brain:** Different subregions of the brain contribute to cognitive function and morphological changes within each region complicates the interpretation of data.
5. **Neural Diversity**: The human brain encompasses an extraordinary variety of neuronal subtypes, glial cells, and specialized subregions. Each cell type may undergo distinct age-related changes (e.g., oxidative stress, proteostatic dysfunction), making it difficult to localize a unifying "cellular/molecular cause" for cognitive impairment.
6. **Inter-individual Variation**: Genetics, epigenetics, lifestyle, and environmental factors vary considerably between individuals, influencing both baseline function and the trajectory of cognitive decline. What underlies decline in one individual may be different from another.
7. **Multifactorial Nature of Cognitive Decline:** Cognitive decline is the result of numerous interacting events (cardiovascular, immune, endocrine, renal, etc.) that compromise brain function.
8. **Comorbid conditions and Convergent Pathologies**: Age-related disorders like cardiovascular disease and metabolic syndrome can affect cerebral blood flow and neuronal health, further complicating the etiology of cognitive impairment. Cognitive decline often results from the interaction of multiple pathological processes; neurodegeneration (e.g., Alzheimer's disease pathology), vascular changes, inflammation, and mitochondrial dysfunction. Untangling which mechanisms initiate the decline in cognitive function within a given individual is inherently difficult.
9. **Lack of Definitive Biomarkers for Early and Subclinical Changes:** Reliable markers that can accurately predict cognitive impairment in the aging population are not currently available.
10. **Imaging and Biochemical Constraints**: While neuroimaging (e.g., MRI, PET) can reveal structural and functional changes within the brain, the techniques do not directly reflect cellular-level processes (e.g. synaptic loss or protein aggregation). Peripheral biomarkers in blood or cerebrospinal fluid show promise but rarely capture the full complexity of brain-specific pathways.
11. **Sensitivity and Specificity**: Biomarkers for Alzheimer's disease (Aβ, tau) or other neuro-degenerative processes are improving, but they are lagging indicators of the subtle cellular changes that may precede cognitive decline.
12. **Slow Progression and Long Timescales:** Cognitive decline occurs over decades, making longitudinal studies expensive and logistically difficult. Capturing early-stage cellular events in a study that spans years is a significant obstacle.
13. **Latency and Compensation**: The brain compensates for neuron loss or dysfunction for extended periods of time, obscuring early indicators of pathophysiology.

### SUPPLEMENT TABLE 2

**Table 2: Barriers to Measuring Cognitive Function in Older Animals**

**1. Health Status and Comorbidities**

a. General Health and Frailty

1. Age-associated pathologies: Older rodents often develop conditions such as kidney dysfunction, chronic inflammation, and neoplasms, which can indirectly affect cognitive performance.
2. Body weight and metabolic issues: Age-related metabolic impairments, including obesity or glucose intolerance, can impact energy balance and endurance during cognitive tasks.

b. Sensory Impairments

1. Vision and hearing loss: Impairments in visual acuity or auditory thresholds may confound tasks like the Morris water maze (which relies on visual cues) or fear conditioning paradigms (which use auditory and visual cues).
2. Olfactory deficits: Olfaction-based tasks, such as odor discrimination, may be affected by impaired olfactory epithelium function.

c. Motor Dysfunction

1. Reduced mobility: Arthritic changes, reduced muscle mass (sarcopenia), and general frailty can impair a rodent's ability to perform tasks requiring active navigation or lever-pressing.
2. Motivation vs. motor deficits: Slowed movement may be misinterpreted as reduced motivation or impaired cognition. Rigorous controls are needed to distinguish between cognitive deficits and compromised motor function.

**2. Behavioral and Environmental Factors**

a. Stress and Anxiety

1. Elevated stress reactivity: Older rodents may have heightened hypothalamic-pituitary-adrenal (HPA) axis activity, influencing performance in stress-sensitive tasks including the elevated plus maze or open field tests.
2. Housing conditions: Enriched versus standard cages can significantly modify both stress resilience and cognitive performance.

b. Circadian Rhythm Changes

1. Altered activity cycles: Aging rodents often exhibit changes in sleep-wake cycles, leading to differences in attention and arousal at various times of the day.
2. Testing Phase: If behavioral testing does not align with the animal's active phase, performance may be confounded by decreased motivation and alertness.

**3. Task-Specific Challenges**

a. Learning Curve and Ceiling Effects

1. Task complexity: Simple tasks may not be sensitive enough to detect subtle age-related deficits, while highly complex tasks can overwhelm older animals.
2. Prior training: Carry-over effects from prior cognitive testing routinely influences subsequent tasks. This is particularly important in longitudinal designs.

b. Motivation and Reward Systems

1. Changes in reward sensitivity: Age-related alterations in dopaminergic and opioid systems can reduce or alter sensitivity to reward, affecting tasks that rely on motivation.
2. Appetitive vs. aversive tasks: Older rodents may respond differently than young animals to aversive vs. appetitive motivation, leading to variable outcomes across paradigms.

**4. Genetic Background and Sex Differences**

a. Strain Variability

1. Strain-specific aging trajectories: Different rodent strains exhibit distinct patterns of neurodegeneration, immune system aging, and lifespan. For example, C57BL/6 mice age differently from BALB/c mice in both behavioral and molecular domains.
2. Mixed vs. inbred strains: Mixed-genotype populations offer higher translational relevance but have increased variability. These models may not be appropriate to isolate the specific mechanisms of cognitive dysfunction with age.

b. Sex-Specific Effects

1. Hormonal fluctuations: Female rodents experience hormonal changes and reproductive senescence that can differentially affect cognition compared to their male counterparts. Reproductive senescence in rodents markedly differs from reproductive senescence in humans.
2. Differential susceptibility to pathologies: Males and females may display different vulnerabilities to neuro-inflammatory changes or oxidative stress with aging.

**5. Additional Limitations of Behavioral Testing**

1. Training and skills of technicians that conduct behavioral tasks are variable and bias is easily incorporated into scientific studies.
2. 'Selection bias' related to the behavioral testing paradigms and the specific dependent variables used to produce the metrics within these paradigms results in poorly validated tests as well as compromised statistical results.
3. Analyses generally include few aged animals and limited trials/animal over several days. In some situations, these comparisons are not sufficient to assess age-related effects and increased trial numbers are necessary to detect differences. Importantly, mice continue to sample/investigate unrewarded locations, and performance occurs in novel environments that introduce stress and anxiety.
4. The number of behavioral tests and the variation in protocols/dependent variables that are being conducted result in testing where one of the fundamental aspects of the scientific method, 'validation of scientific results', cannot be, or is rarely, achieved since the testing is unique to each investigator's laboratory.

#### SUPPLEMENT TABLE 3a

**Table 3a: Variables assessed during the study, abbreviation and description.**

| Behavioral Variable | Abbreviation | Description |
|---|---|---|
| Initial Entries-70% | Ie70 | Entries to reach 70% correct responses based on the trailing 30 trials during acquisition |
| Initial Entries-80% | Ie80 | Entries to reach 80% correct responses based on the trailing 30 trials during acquisition |
| Initial Entries-90% | Ie90 | Entries to reach 90% correct responses based on the trailing 30 trials during acquisition |
| Reversal Entries-70% | Re70 | Entries to reach 70% correct responses based on the trailing 30 trials during the reversal task |
| Reversal Entries-80% | Re80 | Entries to reach 80% correct responses based on the trailing 30 trials during the reversal task |
| Reversal Entries-90% | Re90 | Entries to reach 90% correct responses based on the trailing 30 trials during the reversal task |
| Initial Hours-70% | Ih70 | Time to reach 70% correct responses based on the trailing 30 trials during acquisition |
| Initial Hours-80% | Ih80 | Time to reach 80% correct responses based on the trailing 30 trials during acquisition |
| Initial Hours-90% | Ih90 | Time to reach 90% correct responses based on the trailing 30 trials during acquisition |
| Reversal Hours-70% | Rh70 | Time to reach 70% correct responses based on the trailing 30 trials during the reversal task |
| Reversal Hours-80% | Rh80 | Time to reach 80% correct responses based on the trailing 30 trials during the reversal task |
| Reversal Hours-90% | Rh90 | Time to reach 90% correct responses based on the trailing 30 trials during the reversal task |
| Reversal Learning/Extinction | R-L/E | Increase in correct entries / total entries from hours 51-61 (initial phase of the reversal task) |

### SUPPLEMENT TABLE 3b

**Table 3b:** Model fitting results using Year + Run Date as random effects. *Significance of each variable was based on linear mixed-effects model using Age, Sex, and Age x Sex as fixed effects. Year and Run Date were modeled as random effects. Reversal Measures consistently demonstrated lower variance estimates for Year indicating greater temporal stability and biological sensitivity compared to initial acquisition metrics. p values for rank Re70, Rank Re80 and Rank Re90 indicated that these variables were the most stable metrics across years. Both Rank Rh and Rank R-L/E variables detected gender effects (as well as Rank Ih variables). No Age by Sex interactions were evident. Significant differences in main effects are noted in red text. [1]calculated under 50:50 mixture of a point mass at 0 and $\chi^2_1$ distribution.

| Variable | Age effect p-value | Sex effect p-value | Age × Sex p-value | Estimated variance for random effect 'year' (% of total variance) | p-value[1] for $H_0$: variance for random effect 'year' = 0 |
|---|---|---|---|---|---|
| Rank Ie 70 | 0.0011 | 0.0602 | 0.5912 | 0.0457 (4.6%) | <0.001 |
| Rank Ie 80 | 0.6166 | 0.0005 | 0.7883 | 0.0518 (5.2%) | <0.001 |
| Rank Ie 90 | 0.1283 | <.0001 | 0.4584 | 0.0567 (5.7%) | <0.001 |
| Rank Re 70 | <.0001 | 0.2852 | 0.0919 | 0 (0%) | 1 |
| Rank Re 80 | <.0001 | 0.2580 | 0.2596 | 0 (0%) | 1 |
| Rank Re 90 | <.0001 | 0.8105 | 0.3217 | 0.0034 (0.4%) | 0.240 |
| Rank Ih 70 | <.0001 | 0.0003 | 0.2769 | 0.0856 (8.8%) | <0.001 |
| Rank Ih 80 | 0.0359 | 0.0012 | 0.3965 | 0.0764 (7.7%) | <0.001 |
| Rank Ih 90 | 0.0065 | 0.0039 | 0.4958 | 0.0860 (8.6%) | <0.001 |
| Rank Rh 70 | <.0001 | 0.0038 | 0.5286 | 0.0390 (4.4%) | <0.001 |
| Rank Rh 80 | <.0001 | 0.0033 | 0.5279 | 0.0304 (3.4%) | <0.001 |
| Rank Rh 90 | <.0001 | 0.0434 | 0.3523 | 0.0226 (2.4%) | 0.003 |
| Rank R-L/E | <.0001 | <0.0001 | 0.1853 | 0.0197 (2.5%) | 0.001 |

### SUPPLEMENT TABLE 3b

**Table 3b:** Model fitting results using 'Run Date' or 'Year' as random effects. *Significance of each variable was based on linear mixed-effects model using Age, Sex, and Age x Sex as fixed effects. Run Date and Year were modeled as random effects. Reversal Measures (labeled R) consistently demonstrated lower variance estimates for Year indicating greater temporal stability and biological sensitivity compared to initial acquisition metrics. p values for rank Re70, Rank Re80 and Rank Re90 indicated that these variables were the most stable metrics across years. Both Rank Rh and Rank R-L/E variables detected gender effects (as well as Rank Ih variables). No Age by Sex interactions were evident. Significant differences in main effects are noted in red text. [1]calculated under 50:50 mixture of a point mass at 0 and $\chi_1^2$ distribution.

| Variable | Age effect p-value | Gender effect p-value | Age × gender p-value | Estimated variance for random effect 'run date (% of total variance) | Estimated variance for random effect 'year' (% of total variance) | p-value[1] for $H_0$: variance for random effect 'year' = 0 |
|---|---|---|---|---|---|---|
| Rank Ie70 2 | 0.0011 | 0.0602 | 0.5912 | 0.03584 (3.6%) | 0.04572 (4.6%) | <0.001 |
| Rank Ie 80 2 | 0.6166 | 0.0005 | 0.7883 | 0.02193 (2.2%) | 0.05175 (5.2%) | <0.001 |
| Rank Ie 90 2 | 0.1283 | <.0001 | 0.4584 | 0.02772 (2.8%) | 0.05666 (5.7%) | <0.001 |
| Rank Re 70 2 | <.0001 | 0.2852 | 0.0919 | 0.004053 (0.5%) | 0 (0%) | 1 |
| Rank Re 80 2 | <.0001 | 0.2580 | 0.2596 | 0.005094 (0.6%) | 0 (0%) | 1 |
| Rank Re 90 2 | <.0001 | 0.8105 | 0.3217 | 0.02078 (2.4%) | 0.00338 (0.4%) | 0.240 |
| Rank Ih 70 2 | <.0001 | 0.0003 | 0.2769 | 0.1606 (16.6%) | 0.08555 (8.8%) | <0.001 |
| Rank Ih 80 2 | 0.0359 | 0.0012 | 0.3965 | 0.1474 (14.8%) | 0.0764 (7.7%) | <0.001 |
| Rank Ih 90 2 | 0.0065 | 0.0039 | 0.4958 | 0.1434 (14.3%) | 0.08596 (8.6%) | <0.001 |
| Rank Rh 70 2 | <.0001 | 0.0038 | 0.5286 | 0.01535 (1.7%) | 0.03904 (4.4%) | <0.001 |
| Rank Rh 80 2 | <.0001 | 0.0033 | 0.5279 | 0.02302 (2.6%) | 0.03035 (3.4%) | <0.001 |
| Rank Rh 90 2 | <.0001 | 0.0434 | 0.3523 | 0.02196 (2.4%) | 0.02257 (2.4%) | 0.003 |
| Rank R -L/E | <.0001 | <.0001 | 0.1853 | 0 (0%) | 0.01972 (2.5%) | 0.001 |

### SUPPLEMENT TABLE 4a: STATISTICAL ANALYSIS OF MOVEMENT DURING DARK:LIGHT CYCLES

#### ANALYSIS OF DARK:LIGHT PERIODS

**Dark Phase 1: All Means, se**

| | Mean | se |
|---|---|---|
| 3-4mo | 6.91 | 0.101 |
| 5-6mo | 6.53 | 0.063 |
| 7-10mo | 6.53 | 0.081 |
| 20-22mo | 7.13 | 0.081 |
| 23-25mo | 6.91 | 0.054 |
| 26-30mo | 7.28 | 0.079 |

Data were log-transformed for analysis

**Dark Phase 1: p values**

| | 3-4mo | 5-6mo | 7-10mo | 20-22mo | 23-25mo | 26-30mo |
|---|---|---|---|---|---|---|
| 3-4mo | | 0.019 | 0.037 | 0.534 | 1.0000 | 0.038 |
| 5-6mo | | | 1.0 | <0.001 | <0.001 | <0.001 |
| 7-10mo | | | | <.0001 | 0.0011 | <.0001 |
| 20-22mo | | | | | 0.204 | 0.737 |
| 23-25mo | | | | | | 0.0008 |
| 26-30mo | | | | | | |

**Sex differences:** Female=7.11, se=0.052
Male=6.65, se= 0.038

**Light Phase 1: Female Means, se**

| | Mean | se |
|---|---|---|
| 3-4mo | 3.52 | 0.203 |
| 5-6mo | 3.63 | 0.137 |
| 7-10mo | 4.20 | 0.161 |
| 20-22mo | 4.20 | 0.157 |
| 23-25mo | 4.48 | 0.128 |
| 26-30mo | 4.46 | 0.176 |

Data were log-transformed for analysis

**Light Phase 1: p values**

| | 3-4mo | 5-6mo | 7-10mo | 20-22mo | 23-25mo | 26-30mo |
|---|---|---|---|---|---|---|
| 3-4mo | | 0.998 | 0.096 | 0.086 | 0.009 | 0.007 |
| 5-6mo | | | 0.080 | 0.068 | <0.001 | 0.003 |
| 7-10mo | | | | 1.0 | 0.730 | 0.886 |
| 20-22mo | | | | | 0.736 | 0.891 |
| 23-25mo | | | | | | 1.0 |
| 26-30mo | | | | | | |

**Light Phase 1: Male Means, se**

| | Mean | se |
|---|---|---|
| 3-4mo | 3.55 | 0.156 |
| 5-6mo | 3.23 | 0.090 |
| 7-10mo | 3.91 | 0.124 |
| 20-22mo | 3.76 | 0.128 |
| 23-25mo | 3.74 | 0.074 |
| 26-30mo | 3.45 | 0.113 |

Data were log-transformed

**Light Phase 1: p values**

| | 3-4mo | 5-6mo | 7-10mo | 20-22mo | 23-25mo | 26-30mo |
|---|---|---|---|---|---|---|
| 3-4mo | | 0.474 | 0.455 | 0.910 | 0.875 | 0.995 |
| 5-6mo | | | 0.0001 | 0.010 | 0.0002 | 0.656 |
| 7-10mo | | | | 0.953 | 0.849 | 0.064 |
| 20-22mo | | | | | 1.0 | 0.460 |
| 23-25mo | | | | | | 0.245 |
| 26-30mo | | | | | | |

#### ANALYSIS OF DARK:LIGHT PERIODS

**Dark Phase 2: Female Means, se**

| | Mean | se |
|---|---|---|
| 3-4mo | 7.10 | 0.163 |
| 5-6mo | 6.09 | 0.110 |
| 7-10mo | 6.61 | 0.129 |
| 20-22mo | 6.70 | 0.126 |
| 23-25mo | 6.61 | 0.103 |
| 26-30mo | 7.10 | 0.141 |

Data were log-transformed for analysis

**Dark Phase 2: p values**

| | 3-4mo | 5-6mo | 7-10mo | 20-22mo | 23-25mo | 26-30mo |
|---|---|---|---|---|---|---|
| 3-4mo | | <0.0001 | 0.172 | 0.390 | 0.107 | 1.000 |
| 5-6mo | | | 0.026 | 0.003 | 0.008 | <0.0001 |
| 7-10mo | | | | 0.995 | 1.000 | 0.109 |
| 20-22mo | | | | | 0.991 | 0.297 |
| 23-25mo | | | | | | 0.055 |
| 26-30mo | | | | | | |

**Dark Phase 2: Male Means, se**

| | Mean | se |
|---|---|---|
| 3-4mo | 6.79 | 0.125 |
| 5-6mo | 6.70 | 0.073 |
| 7-10mo | 6.62 | 0.100 |
| 20-22mo | 6.74 | 0.103 |
| 23-25mo | 6.30 | 0.059 |
| 26-30mo | 6.61 | 0.091 |

Data were log-transformed for analysis

**Dark Phase 2: p values**

| | 3-4mo | 5-6mo | 7-10mo | 20-22mo | 23-25mo | 26-30mo |
|---|---|---|---|---|---|---|
| 3-4mo | | 0.986 | 0.883 | 0.100 | 0.005 | 0.840 |
| 5-6mo | | | 0.987 | 0.999 | <0.001 | 0.973 |
| 7-10mo | | | | 0.958 | 0.068 | 1.000 |
| 20-22mo | | | | | 0.003 | 0.932 |
| 23-25mo | | | | | | 0.051 |
| 26-30mo | | | | | | |

**Light Phase 2: Female Means, se**

| | Mean | se |
|---|---|---|
| 3-4mo | 3.78 | 0.195 |
| 5-6mo | 3.86 | 0.132 |
| 7-10mo | 4.27 | 0.154 |
| 20-22mo | 4.33 | 0.151 |
| 23-25mo | 4.55 | 0.123 |
| 26-30mo | 4.72 | 0.169 |

Data were log-transformed for analysis

**Light Phase 2: p values**

| | 3-4mo | 5-6mo | 7-10mo | 20-22mo | 23-25mo | 26-30mo |
|---|---|---|---|---|---|---|
| 3-4mo | | 0.999 | 0.362 | 0.232 | 0.012 | 0.004 |
| 5-6mo | | | 0.335 | 0.186 | 0.002 | 0.001 |
| 7-10mo | | | | 0.999 | 0.727 | 0.371 |
| 20-22mo | | | | | 0.870 | 0.518 |
| 23-25mo | | | | | | 0.965 |
| 26-30mo | | | | | | |

**Light Phase 2: Male Means, se**

| | Mean | se |
|---|---|---|
| 3-4mo | 3.83 | 0.149 |
| 5-6mo | 3.56 | 0.087 |
| 7-10mo | 3.92 | 0.119 |
| 20-22mo | 3.97 | 0.123 |
| 23-25mo | 4.02 | 0.071 |
| 26-30mo | 3.34 | 0.109 |

**Light Phase 2: p values**

| | 3-4mo | 5-6mo | 7-10mo | 20-22mo | 23-25mo | 26-30mo |
|---|---|---|---|---|---|---|
| 3-4mo | | 0.641 | 0.997 | 0.977 | 0.857 | 0.086 |
| 5-6mo | | | 0.149 | 0.073 | 0.001 | 0.591 |
| 7-10mo | | | | 0.999 | 0.980 | 0.004 |
| 20-22mo | | | | | 0.999 | 0.002 |
| 23-25mo | | | | | | <0.001 |
| 26-30mo | | | | | | |

### TABLE 4b: STATISTICAL ANALYSIS OF MOVEMENT DURING DARK:LIGHT CYCLES

#### ANALYSIS OF DARK:LIGHT PERIODS

**Dark Phase 3: Female Means, se**

| | Mean | se |
|---|---|---|
| 3-4mo | 7.10 | 0.197 |
| 5-6mo | 6.15 | 0.134 |
| 7-10mo | 6.21 | 0.156 |
| 20-22mo | 6.45 | 0.153 |
| 23-25mo | 6.5 | 0.124 |
| 26-30mo | 7.2 | 0.171 |

Data were log-transformed for analysis

**Dark Phase 3: p values**

| | 3-4mo | 5-6mo | 7-10mo | 20-22mo | 23-25mo | 26-30mo |
|---|---|---|---|---|---|---|
| 3-4mo | | 0.001 | 0.005 | 0.093 | 0.105 | 0.999 |
| 5-6mo | | | 0.999 | 0.683 | 0.383 | <0.001 |
| 7-10mo | | | | 0.879 | 0.676 | 0.001 |
| 20-22mo | | | | | 0.999 | 0.015 |
| 23-25mo | | | | | | 0.014 |
| 26-30mo | | | | | | |

**Dark Phase 3: Male Means, se**

| | Mean | se |
|---|---|---|
| 3-4mo | 7.04 | 0.151 |
| 5-6mo | 6.54 | 0.088 |
| 7-10mo | 6.12 | 0.121 |
| 20-22mo | 6.33 | 0.124 |
| 23-25mo | 6.27 | 0.072 |
| 26-30mo | 6.62 | 0.110 |

Data were log-transformed for analysis

**Dark Phase 3: p values**

| | 3-4mo | 5-6mo | 7-10mo | 20-22mo | 23-25mo | 26-30mo |
|---|---|---|---|---|---|---|
| 3-4mo | | 0.049 | <0.001 | 0.004 | <0.001 | 0.213 |
| 5-6mo | | | 0.059 | 0.724 | 0.148 | 0.994 |
| 7-10mo | | | | 0.852 | 0.914 | 0.030 |
| 20-22mo | | | | | 0.998 | 0.494 |
| 23-25mo | | | | | | 0.078 |
| 26-30mo | | | | | | |

**Light Phase 3: F and M Means, se**

| | Mean | se |
|---|---|---|
| 3-4mo | 3.88 | 0.144 |
| 5-6mo | 3.89 | 0.090 |
| 7-10mo | 4.27 | 0.115 |
| 20-22mo | 4.35 | 0.116 |
| 23-25mo | 4.74 | 0.077 |
| 26-30mo | 4.13 | 0.112 |

Data were log-transformed for analysis

**Light Phase 3: p values**

| | 3-4mo | 5-6mo | 7-10mo | 20-22mo | 23-25mo | 26-30mo |
|---|---|---|---|---|---|---|
| 3-4mo | | 1.00 | 0.269 | 0.111 | <0.001 | 0.731 |
| 5-6mo | | | 0.083 | 0.019 | <0.001 | 0.509 |
| 7-10mo | | | | 0.997 | 0.007 | 0.953 |
| 20-22mo | | | | | 0.044 | 0.758 |
| 23-25mo | | | | | | <0.001 |
| 26-30mo | | | | | | |

#### SUPPLEMENT TABLE 5. Selection Criteria for Genes based on Inferential Analysis (Partial Spearman)

| CATEGORY | Criteria 1 | Criteria 2 | Criteria 3 |
|---|---|---|---|
| Cognition Associated | Partial Spearman (Pseudotime \| Age) $p<0.05$ | Partial Spearman (Age \| Pseudotime) $p\geq 0.05$ | Partial Spearman (Pseudotime \| Age) $\lvert\rho\rvert\geq 0.44$ |
| Shared Age & Cognition Associated | Partial Spearman (Pseudotime \| Age) $p<0.05$ | Partial Spearman (Age \| Pseudotime) $p<0.05$ | Partial Spearman (Pseudotime \| Age) $\lvert\rho\rvert\geq 0.44$ (Age \| Pseudotime) $\lvert\rho\rvert\geq 0.44$ |
| Age Associated | Partial Spearman (Age \| Pseudotime) $p<0.05$ | Partial Spearman (Pseudotime \| Age) $p\geq 0.05$ | Partial Spearman (Age \| Pseudotime) $\lvert\rho\rvert\geq 0.44$ |

#### SUPPLEMENT Figure 1

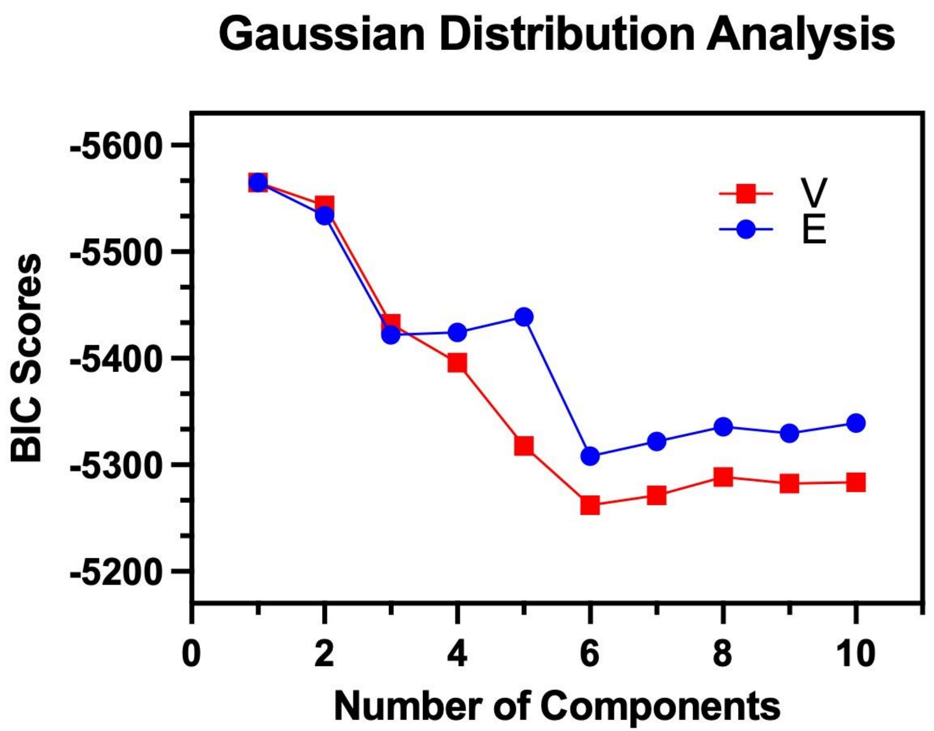


**Supplement Figure 1:** Bayesian information criterion (BIC) scores were used to determine the optimal number of components in Gaussian mixture models applied to the dataset. Models with either variable (V) or equal (E) variance across components were evaluated across 1–10 components. Lower BIC values indicate improved model fit. Both variance structures show a marked improvement in fit up to four components, with diminishing returns or plateauing thereafter. Based on these results, a four-component model was selected as the optimal representation of the underlying data structure.

#### SUPPLEMENT Figure 2

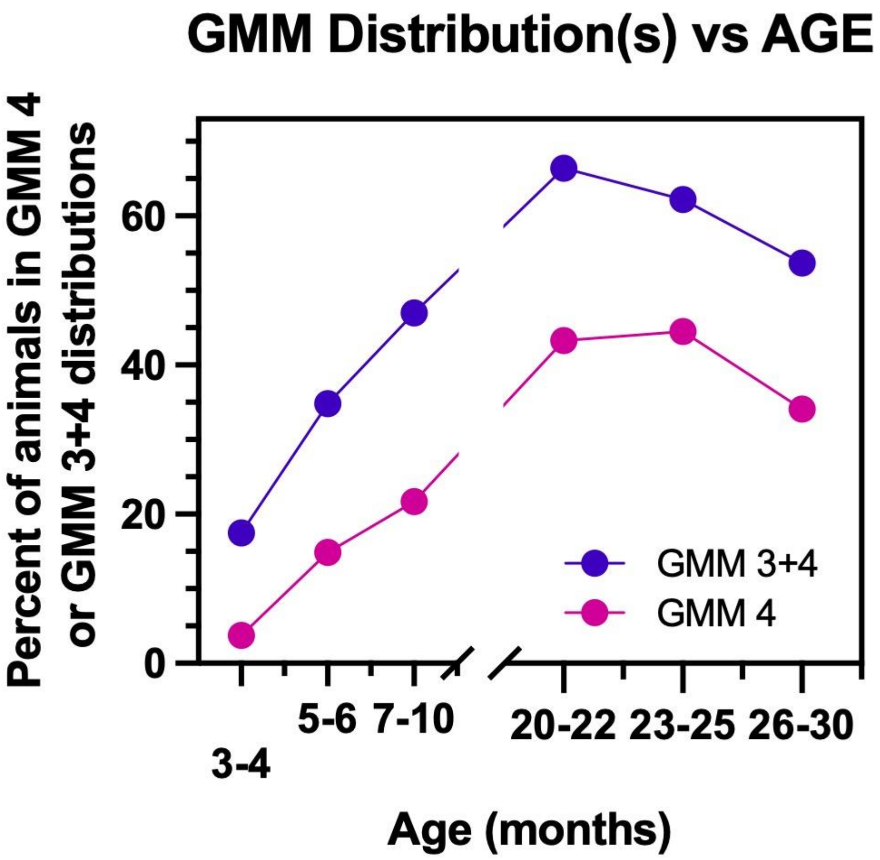


**Supplement Figure 2. Age-dependent increase in animals assigned to higher Gaussian mixture model (GMM) components** The percentage of animals assigned to higher-order Gaussian mixture model components is shown across age groups. Animals classified within GMM component 4 (magenta) and combined components 3 and 4 (purple) increase progressively with age indicating an age-associated shift toward impaired cognitive performance. This trend is evident from early adulthood (5-6mo) through advanced age, with the highest proportions observed in older cohorts. These findings demonstrate that increasing age is associated with an increasing probability of cognitive impairment.

### SUPPLEMENTAL FIGURE 3. DESCRIPTIVE ANALYSIS

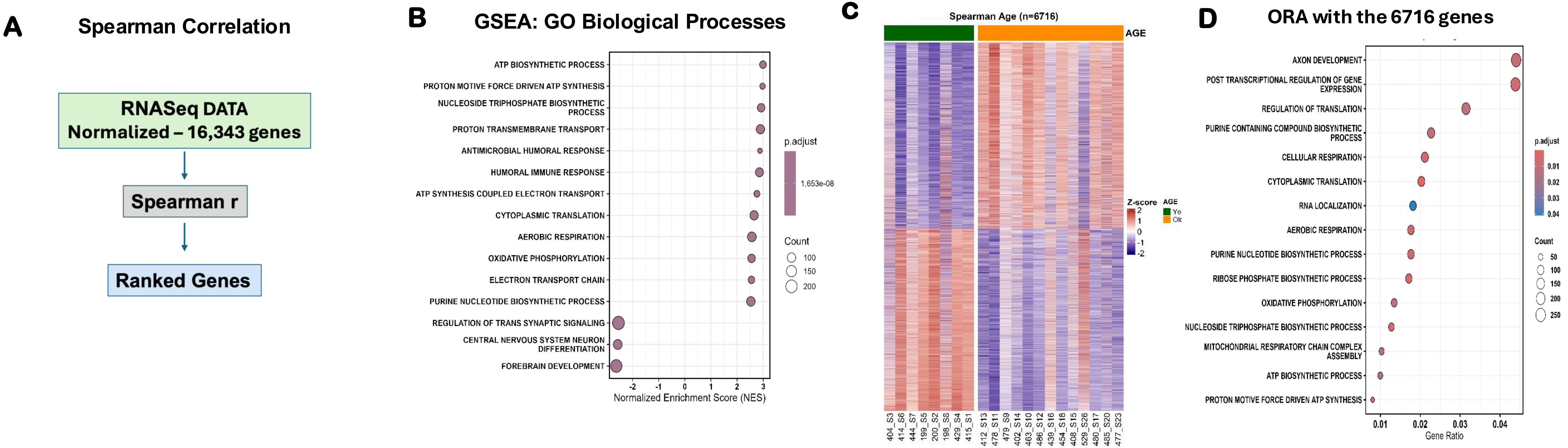


**Supplemental Figure 3. Descriptive analysis captures broad age-associated transcriptional programs**

(A) Schematic overview of the descriptive analysis workflow. Normalized RNA-seq data (16,343 genes) were analyzed using bivariate Spearman correlations to identify genes associated with age without accounting for shared variance with cognitive performance. (B) Gene set enrichment analysis (GSEA) of genes ranked by Spearman correlation with age. Enriched pathways are dominated by metabolic, mitochondrial, and translational processes, reflecting broad age-associated transcriptional changes. (C) Heatmap of age-associated genes (n = 6,716) showing separation between young and old animals, consistent with widespread transcriptional remodeling across aging. (D) Over-representation analysis (ORA) of age-associated genes further highlights enrichment of canonical aging pathways, including oxidative phosphorylation, biosynthetic processes, and cellular metabolism. Together, these analyses demonstrate that descriptive (bivariate) approaches capture global age-related transcriptional changes but lack specificity for cognition-associated pathways, contrasting with the cilium- and microtubule-related programs identified using inferential (variance-partitioned) analyses in the main manuscript.

### Supplemental Figure 4

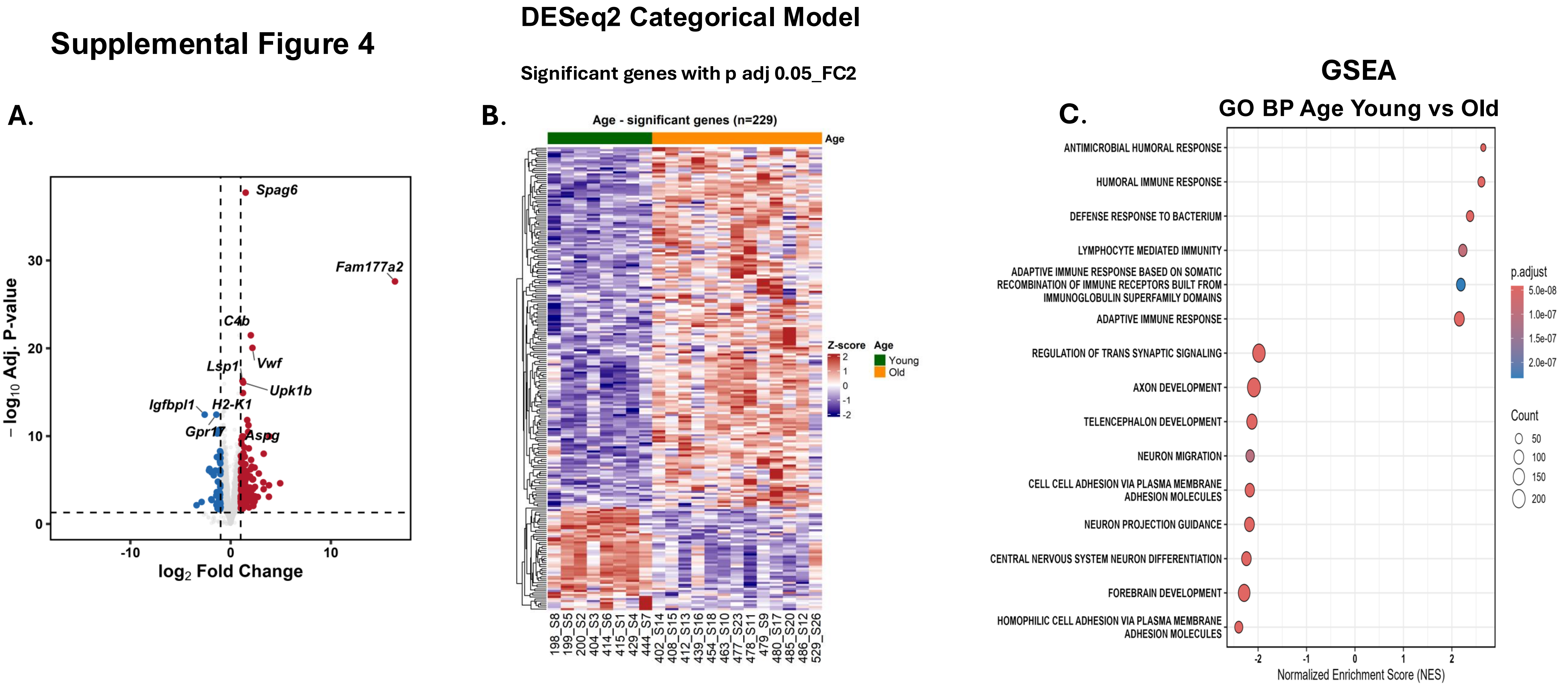


**Supplemental Figure 4. Age-based differential expression captures canonical aging pathways but not cognition-associated programs**
(A) Volcano plot of differential gene expression between young and old animals using a categorical DESeq2 model. Genes meeting significance thresholds (adjusted $p < 0.05$, $|\log_2$ fold change$| \geq 1$) are highlighted, illustrating transcripts upregulated or downregulated with age. (B) Heatmap of significantly differentially expressed genes ($n = 229$) across individual animals, showing clear separation between young and old groups and reflecting widespread age-associated transcriptional changes. (C) Gene Ontology (GO) biological process enrichment analysis of age-associated genes reveals overrepresentation of canonical aging pathways, including immune and inflammatory responses and neuronal development–related processes. Together, these analyses demonstrate that standard age-group differential expression captures broad aging-associated transcriptional changes but does not identify the cilium- and microtubule-associated programs linked to cognitive performance revealed by trajectory-based and partial correlation analyses.

### Supplemental Figure 5

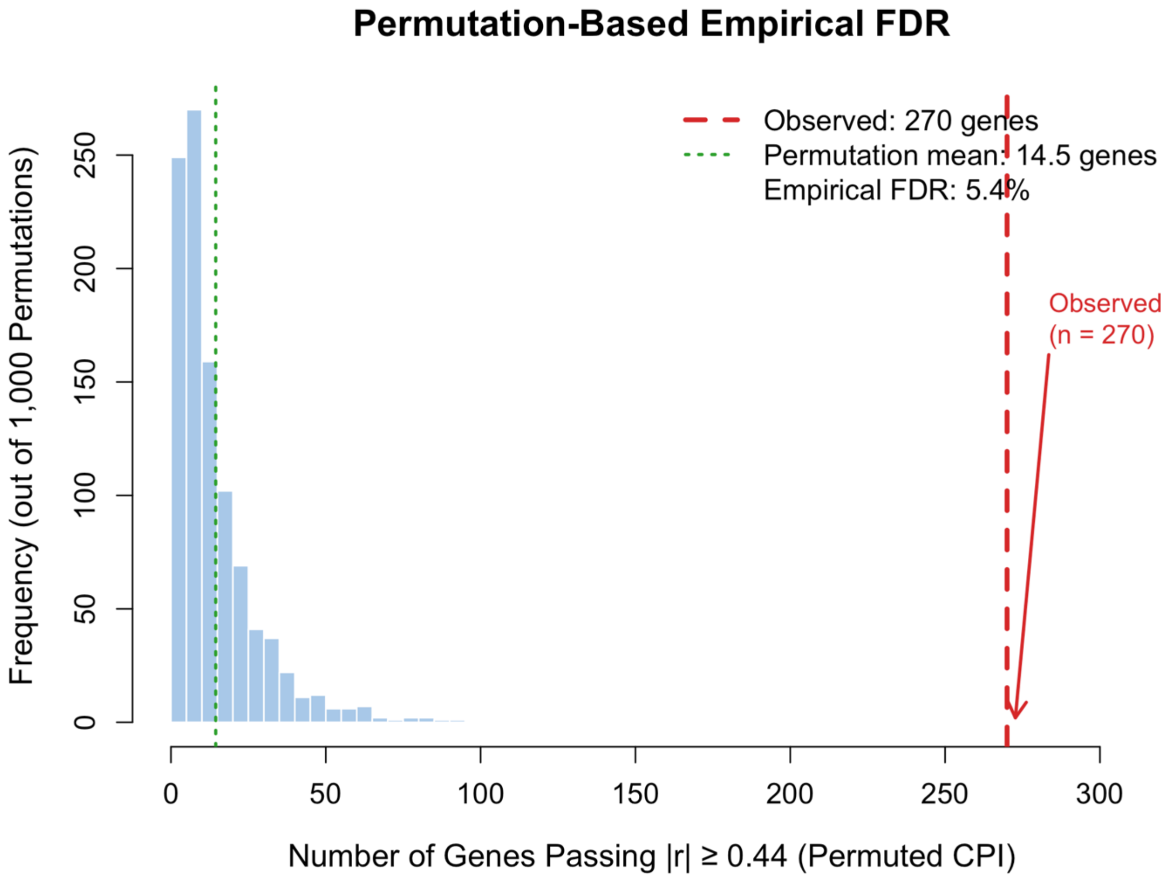


**Supplementary Figure 5. Permutation-based empirical false discovery rate analysis confirms that CPI-correlated hippocampal genes exceed chance expectations.** Histogram shows the null distribution of gene counts generated by randomly shuffling Cognitive Performance Index (CPI) values across the 21 animals and recomputing partial Spearman correlations (controlling for age) for all candidate genes across 1,000 independent permutations (blue bars). The green dotted line indicates the permutation mean (14.5 genes), representing the number of genes expected to exceed $|r| \geq 0.44$ by chance alone. The red dashed line indicates the observed gene count (n = 271), which is approximately 19-fold greater than the permutation mean and falls entirely outside the null distribution. No single permutation approached the observed count. Empirical false discovery rate = 5.4%, indicating that approximately 94.6% of the 271 identified genes reflect genuine associations between hippocampal gene expression and cognitive performance independent of chronological age. CPI, Cognitive Performance Index; FDR, false discovery rate.

### SUPPLEMENTAL FIGURE 6. Redundancy analysis of cilia- and microtubule-related gene sets

#### A. Number of cilia related pathways: 18

Cilia associated terms: CILIA,CILIUM,AXONEME,FLAGELLUM,MICROTUBULE

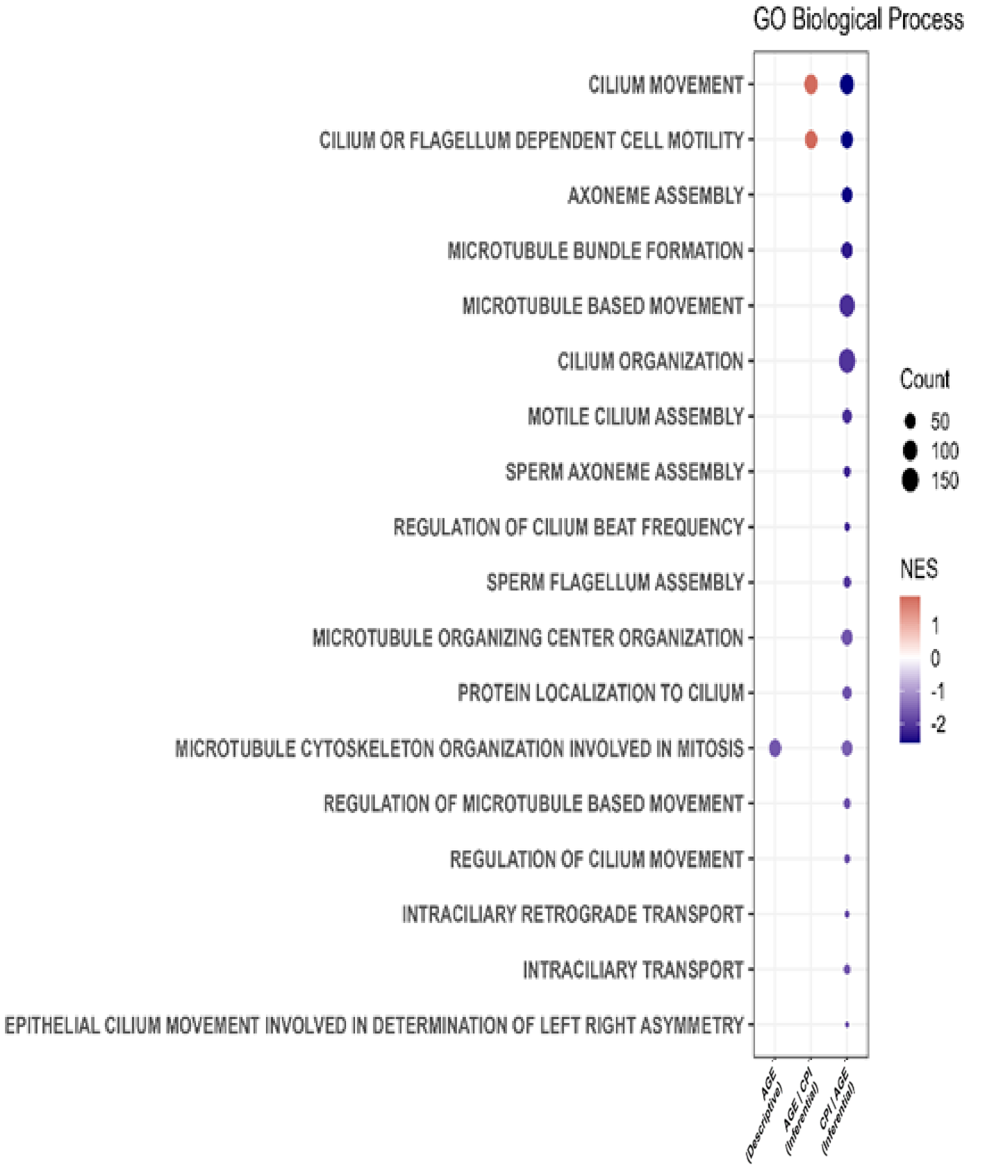


#### B. Redundancy Summary:

**Total number of leading-edge genes: 307**

**152 shared and 155 unique genes**

| Number of Pathways (N) | Gene Count | Percent of Total Genes (307) |
|---|---|---|
| 1 | 155 | 50.49% |
| 2 | 45 | 14.66% |
| 3 | 42 | 13.68% |
| 4 | 12 | 3.91% |
| 5 | 13 | 4.23% |
| 6 | 13 | 4.23% |
| 7 | 5 | 1.63% |
| 8 | 4 | 1.30% |
| 9 | 14 | 4.56% |
| 10 | 1 | 0.33% |
| 12 | 3 | 0.98% |

#### C. Representative Ciliary/Microtubule Biological Processes: 8

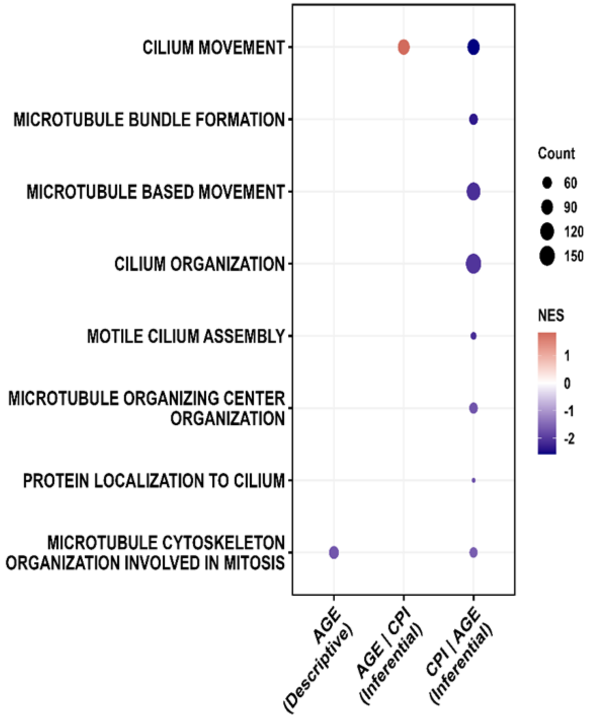


## D.

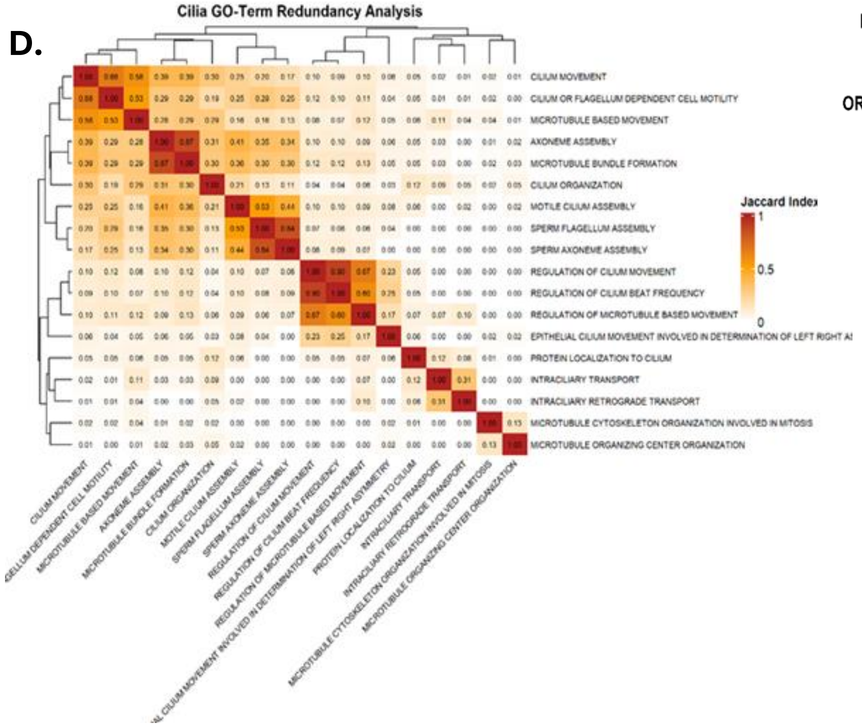

**SUPPLEMENTAL FIGURE 6. Redundancy analysis of cilia- and microtubule-related gene sets**

(A) Gene Ontology (GO) biological process enrichment for cilia- and microtubule-related pathways identified in the CPI | AGE model. Eighteen significantly enriched pathways associated with ciliary structure, axonemal assembly, and microtubule-based processes were identified using keyword filtering (cilia, axoneme, flagellum, microtubule). Enrichment was minimal or absent in descriptive Age and AGE | CPI models, indicating specificity to cognition-associated gene sets. (B) Summary of leading-edge gene overlap across the 18 enriched pathways. A total of 307 genes were identified, with approximately half unique to individual pathways, indicating that enrichment reflects a distributed transcriptional program rather than a small set of shared genes. (C) Representative non-redundant biological processes after removal of pathways with complete gene overlap. Eight pathways capturing distinct aspects of ciliary and microtubule biology were retained, illustrating the diversity of the underlying molecular program associated with cognitive performance. (D) Pairwise pathway similarity assessed using the Jaccard index. Clustering reveals groups of related pathways with partial overlap, supporting the presence of structured redundancy while confirming that the retained pathways represent distinct biological processes. Together, these analyses demonstrate that enrichment of cilia- and microtubule-related pathways in the CPI | AGE model. reflects a robust, distributed, and biologically meaningful transcriptional program rather than redundancy-driven enrichment

#### SUPPLEMENTAL FIGURE 7.

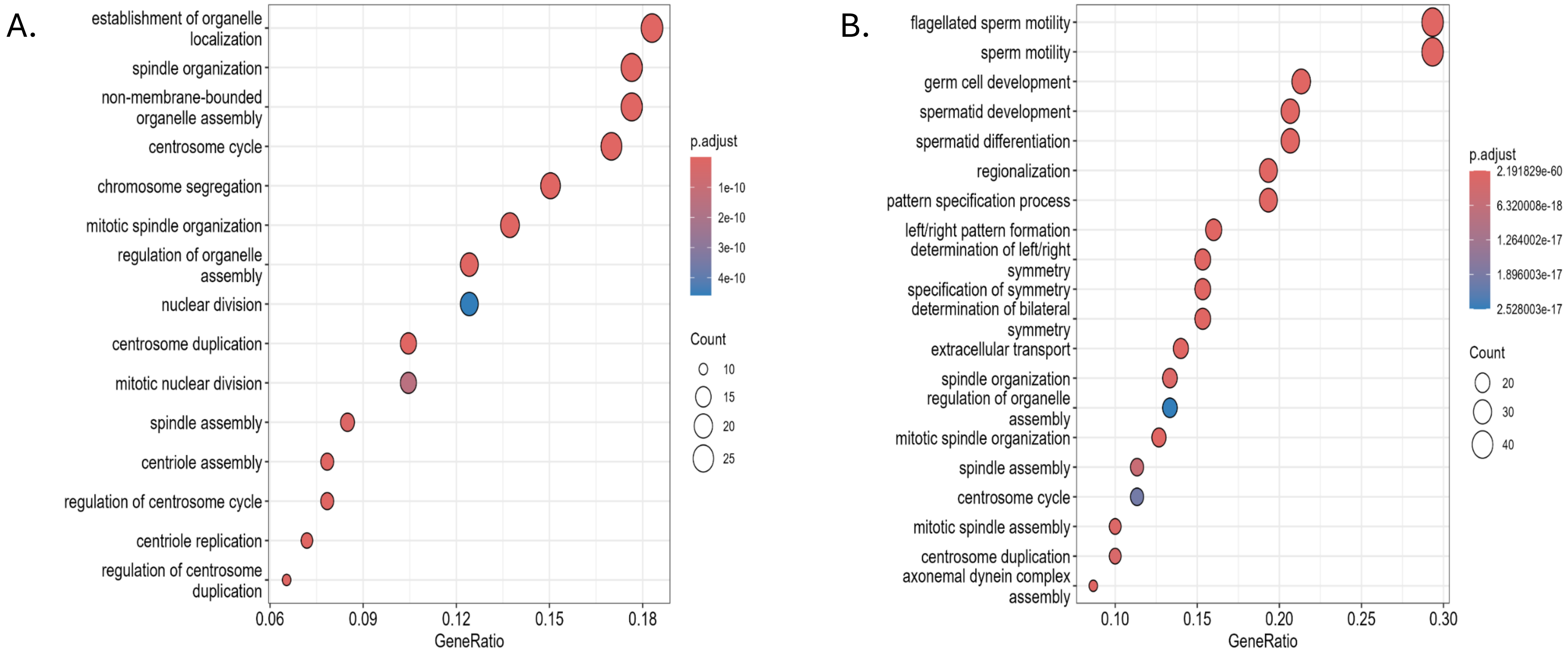


**Supplementary Figure 7. Analysis of non-ciliary pathway enrichment among cilia-associated genes**
Evaluation of potential non-ciliary biological pathways represented within the gene set identified through cilia-focused screening. A total of 307 genes were examined, comprising 155 unique genes and 152 genes shared across multiple cilia-related pathways. Enrichment analysis of these genes revealed additional biological processes beyond canonical ciliary function, including pathways related to cellular metabolism, signaling, and stress responses. Despite the presence of these secondary associations, the majority of genes were initially identified through cilia- and microtubule-related pathways, indicating that non-ciliary enrichments reflect broader functional integration rather than independent pathway dominance. These findings support the specificity of the cilia-associated signal while highlighting the interconnected nature of ciliary biology with other cellular processes.

### Supplemental Figure 8

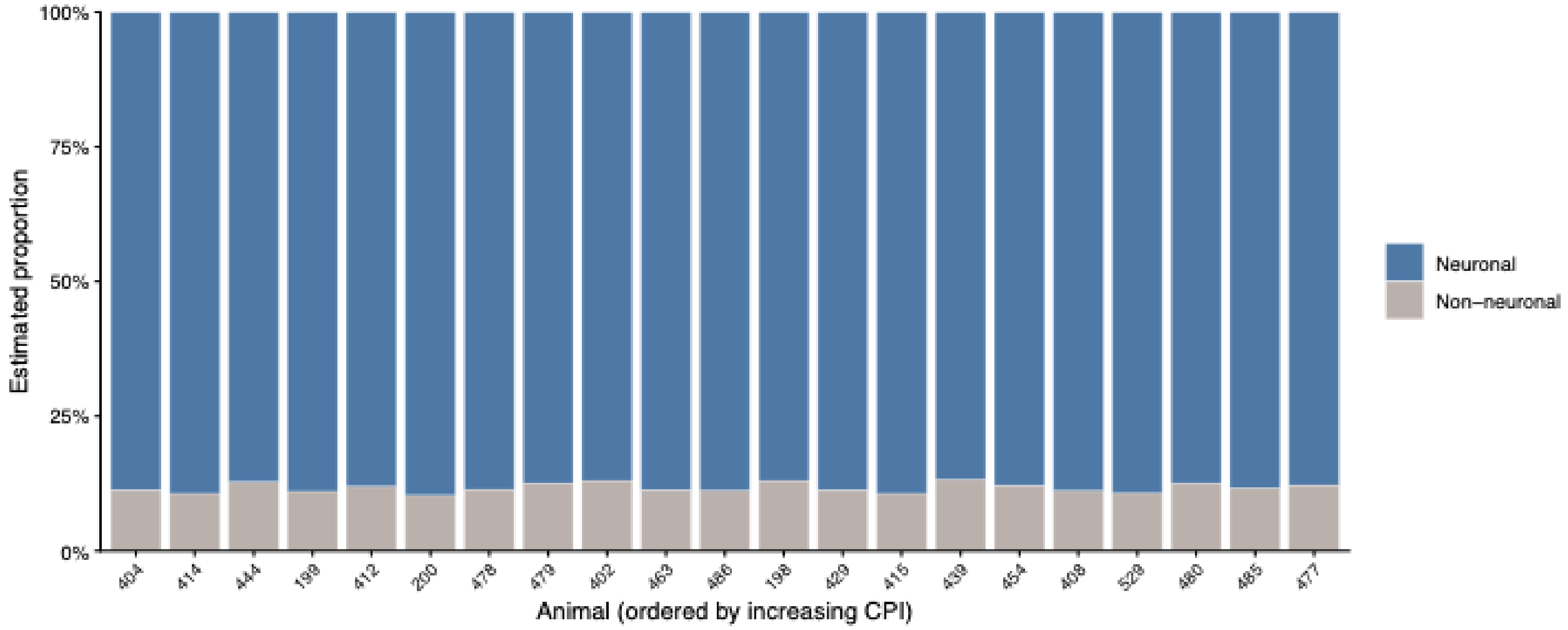


**Supplemental Figure 8.** The estimated hippocampal cellular composition is predominantly neuronal across all animals regardless of cognitive performance. Estimated cell-type proportions from Figure x collapsed into neuronal and non-neuronal compartments, shown for each of 21 animals ordered left to right by increasing CPI. Neuronal cell types (Cajal-Retzius, GABA-Lhx6, granule-immature, granule-mature, Mossy-Cyp26b1, Neuroblast_1, Neuroblast_2, and nIPC) are distinguished from non-neuronal types (astrocytes, endothelial cells, microglia, oligodendrocytes, OPCs, and radial glia-like cells). The neuronal compartment constitutes the large majority of the estimated composition in every animal and shows no systematic shift across the CPI gradient, indicating that the cognition-associated gene expression signature is not attributable to a change in the relative abundance of neuronal versus non-neuronal cells. n = 21 animals.

### Supplemental Figure 9

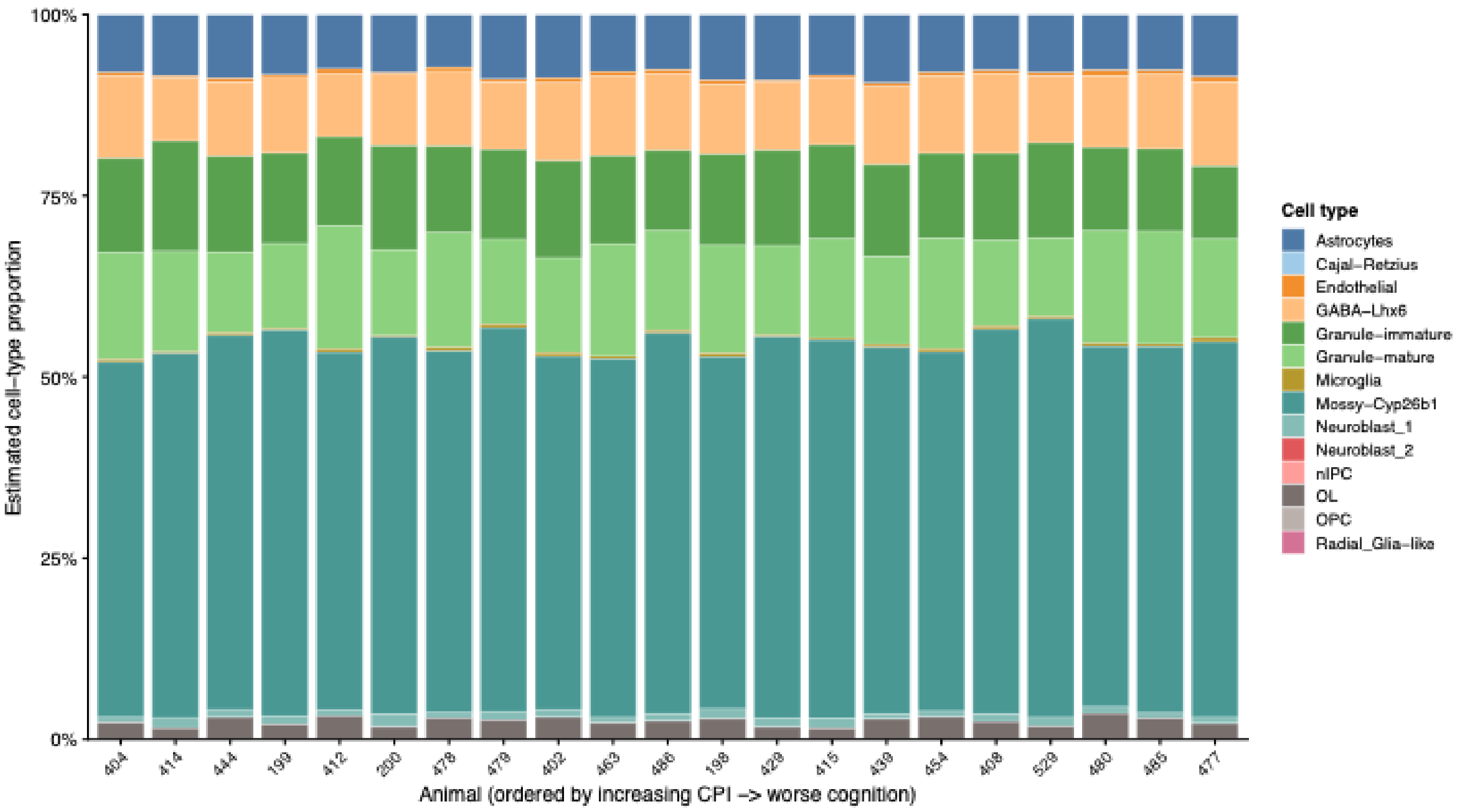


**Supplemental Figure 9. CIBERSORTx deconvolution resolves 14 hippocampal cell types across animals spanning the full range of cognitive performance.**

Estimated relative proportions of 14 hippocampal cell types, derived by CIBERSORTx deconvolution of bulk hippocampal RNA-seq using a single-cell reference (Zeisel et al., 2018), shown for each of 21 animals. Animals (columns) are ordered left to right by increasing Cognitive Performance Index (CPI; higher values indicate greater cognitive impairment). Each colored segment represents the fractional contribution of one cell type to the estimated cellular composition, normalized to 100% per animal. Neuronal populations, predominantly Mossy-Cyp26b1 neurons and mature and immature granule cells, account for the majority of the estimated composition across all animals. Cell-type proportions are broadly stable across the CPI gradient, consistent with the absence of a significant association between individual cell-type proportions and the cilia structural module score after age correction (all $p > 0.18$; Table Z). $n = 21$ animals.

### Supplemental Figure 10

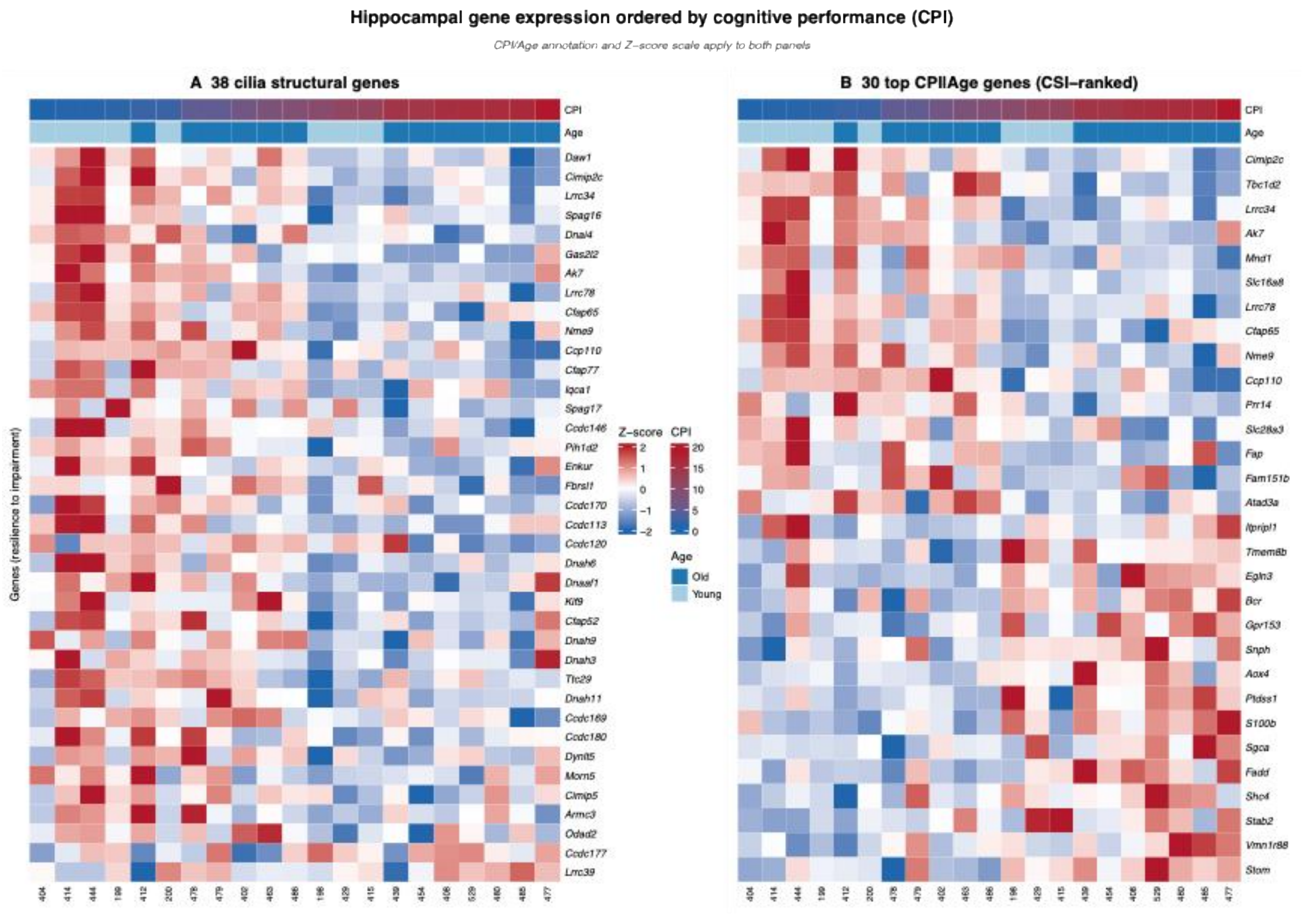


**Supplemental Figure 10. Cilia structural genes are uniformly associated with cognitive resilience, in contrast to the functionally heterogeneous top-ranked gene set.** Hippocampal gene expression heatmaps for 21 aged and young mice, with animals (columns) ordered left to right by increasing Cognitive Performance Index (CPI; higher values indicate greater cognitive impairment). Expression values are VST-normalized and Z-scored by gene (row); red indicates higher and blue lower relative expression. The CPI gradient and age group (young, 5–6 months; old, 21–25 months) are annotated above each panel. Genes (rows) are ordered by their Spearman correlation with CPI, from most resilience-associated (top) to most impairment-associated (bottom). (A) The 38 bona fide cilia structural genes within the CPI|Age set. Expression decreases consistently with worsening cognition across nearly all genes, with only Lrrc39 and Ccdc177 showing the opposite pattern, reflecting the uniform resilience association of cilia structural genes (36 of 38 genes; consistent sign consistency across leave-one-out resampling). (B) The 30 top-ranked CPI|Age genes by Cognitive Stability Index. In contrast to panel A, this functionally heterogeneous set displays a bidirectional pattern: resilience-associated genes (including the seven cilia genes shared with panel A) cluster at the top, whereas impairment-associated immune and metabolic genes (e.g., Stom, S100b, Fadd, Vmn1r88) occupy the lower rows. The CPI/Age annotation and Z-score scale apply to both panels. n = 21 animals.

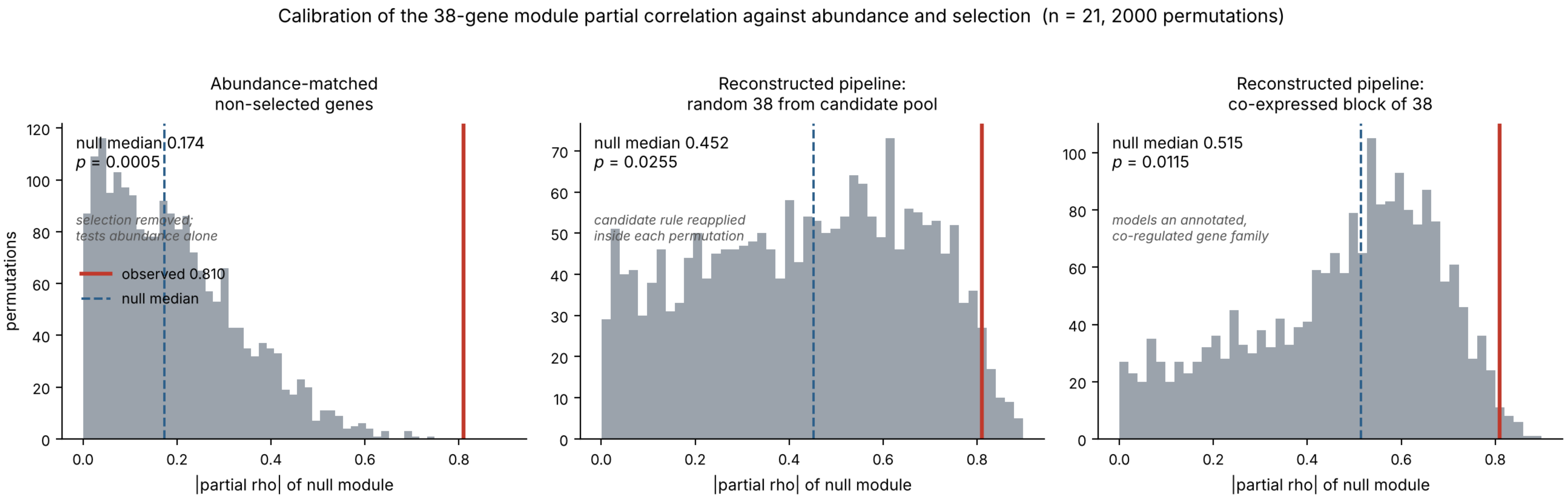


**Supplementary Figure 11. Calibration of the 38-gene cilia structural module against transcript abundance and gene selection.** The module-level partial Spearman correlation with the cognitive performance index, controlling for age (observed |rho| = 0.810, red line), was evaluated against three null distributions, each generated from 2,000 permutations (n = 21 animals; blue dashed line, null median). **(Left)** Abundance control. Gene sets of 38 were drawn from genes not subject to selection, matched gene by gene to the mean variance-stabilized expression of the 38 module genes, and scored identically. Although the module genes lie in the lowest decile of the expression distribution (median 11th percentile), the observed value exceeded all 2,000 abundance-matched sets (null median 0.174, empirical p = 0.0005), establishing that the association is not a consequence of low transcript abundance. **(Center)** Selection control. The cognitive performance index was permuted across animals with age held to its original animal, the candidate-selection criteria were reapplied within each permutation, and 38 genes were drawn at random from the resulting candidate pool, modelling the cilia annotation as a criterion independent of cognitive performance. The observed value was exceeded in 50 of 2,000 permutations (null median 0.452, empirical p = 0.026). **(Right)** Same as center, but with the 38 genes drawn as a mutually co-expressed block within the candidate pool, modelling the module as a coherent, co-regulated gene family; the observed value was exceeded in 22 of 2,000 permutations (null median 0.515, empirical p = 0.012). Because the module genes were selected on their association with the cognitive performance index in this same cohort, approximately half of the observed magnitude is reproducible from the selection procedure alone, and the nominal p value substantially overstates the strength of the association. The association nonetheless exceeds the calibrated null under both selection models. The more conservative value (p = 0.026) is reported in the Abstract and Results.
